# Sediments from a seasonally euxinic coastal ecosystem show high nitrogen cycling potential

**DOI:** 10.1101/2025.02.26.640368

**Authors:** Isabel M. L. Rigutto, Ştefania C. Sburlan, Lars W. P. de Bont, Tom Berben, Rob M. de Graaf, Caroline P. Slomp, Mike S. M. Jetten

**Affiliations:** Department of Microbiology, Radboud Institute for Biological and Environmental Sciences, Radboud University, Nijmegen, the Netherlands; Department of Earth Sciences - Geochemistry, Utrecht University, Utrecht, the Netherlands

## Abstract

Coastal ecosystems are susceptible to eutrophication and deoxygenation, which may alter their nitrogen cycle dynamics. Here, we investigated the microbial nitrogen cycling potential in the sediment of a seasonally euxinic coastal ecosystem (Lake Grevelingen, NL), in winter and summer. Porewater profiles showed ammonium (NH_4_^+^) concentrations exceeding 10 mM and rapid depletion of electron acceptors with depth. Activity tests revealed NH_4_^+^ oxidation potential up to 53 µmol g^−1^ day^−1^, even in anoxic sediment layers. A nitrifying microbial community was present in both oxic and anoxic sediment sections (up to 1.4% relative abundance). NO₃^⁻^, nitrite (NO_2_^−^) and nitrous oxide (N_2_O) reduction potential were prominent across all sediment sections, with the highest rates (167 µmol NO_3_^−^ g^−1^ day^−1^) in the surface sediment in summer. Denitrification (79.3-98.4%) and dissimilatory nitrate reduction to ammonium (DNRA; 1.6-20.7%) were the major NO_3_^−^ removal pathways, as supported by the detection of the *narG/napA, nirK/nirS, norB, nosZ* and *nrfA/otr* genes in all sediment sections. The DNRA contribution increased with depth and with the addition of electron donors, such as monomethylamine. Anaerobic ammonium oxidation (anammox) was not detected in these eutrophic sediments. Combined, our results show that there is high potential for nitrogen removal in eutrophic coastal ecosystems which may help further restoration measures.

## Introduction

Coastal ecosystems connect land to the open sea, rendering them an important filter for the nutrients that enter marine waters from terrestrial ecosystems (Kuliński et al., 2022). Yet, the biogeochemical functioning of this coastal filter is strongly impacted by shifts in environmental conditions due to anthropogenic activities. Intensive agricultural fertilizer use, husbandry, and industrial emissions may lead to large discharges of nutrients into rivers, resulting in an increased influx of nutrients into coastal systems (Loughner et al., 2016; Lu et al., 2018; Seitzinger et al., 2010). These excess nutrient loads stimulate productivity and the subsequent degradation of organic matter through aerobic respiration, consuming (nearly all) oxygen (Breitburg et al., 2018; Kessouri et al., 2021). This leads to severe deoxygenation, which is further enhanced by warming of the coastal ocean and seas (Breitburg et al., 2018; Schmidtko et al., 2017; Wei et al., 2021). Upon increased temperatures, the oxygen solubility in marine waters declines, and stratification in these waters intensifies, which hampers the transport of oxygen between the different water layers (Oschlies et al., 2018). Enhanced eutrophication and deoxygenation can push nutrient cycles out of balance. Nitrogen is an essential and often limiting nutrient, and therefore alterations in the type and amount of fixed nitrogen may affect marine productivity directly. Moreover, shifts in the marine nitrogen cycle can affect the production of the very potent greenhouse gas nitrous oxide (N_2_O), as well as other greenhouse gases such as carbon dioxide (CO_2_) and methane (CH_4_), through the tight link between the nitrogen and carbon cycles (Long et al., 2021).

The marine nitrogen cycle is the result of the complex interplay between the various microbial guilds that can either cooperate or compete for resources (Kuypers et al., 2018). Sinking organic matter is mineralized in marine bottom waters and sediments, where the formed ammonium (NH_4_^+^) can be used in various microbial processes. In nitrification, NH_4_^+^ is oxidized to nitrite (NO_2_^−^) and subsequently to nitrate (NO_3_^−^) with oxygen as electron acceptor. NH_4_^+^ oxidation is catalyzed by ammonia-oxidizing bacteria (AOB) and ammonia-oxidizing archaea (AOA) that contain *amoCAB* and *hao* genes (Francis et al., 2005; Klotz et al., 1997; Sayavedra-Soto et al., 1994). AOB belonging to the genera *Nitrosomonas, Nitrospira* and *Nitrosococcus* have been detected in marine environments (Liu, Cheng, et al., 2023), and thaumarchaeal AOA can be very abundant in oxic marine waters (Wuchter et al., 2006). Nitrite-oxidizing bacteria (NOB) possess *nxrAB* genes and are phylogenetically diverse with *Nitrospina* being considered the most abundant marine NOB (Beman et al., 2013; Sun et al., 2019). Complete ammonia-oxidizing bacteria (comammox) belong to the *Nitrospira* genus (Daims et al., 2015; Van Kessel et al., 2015). Comammox bacteria have been detected in tidal flats (Sun et al., 2020), continental shelves (Liu, Xu, et al., 2023) and salt marshes (Bernhard et al., 2021) by qPCR-based approaches, but not yet in the open ocean. Although NH_4_^+^ oxidation requires oxygen, nitrification at nanomolar oxygen levels has been observed (Bristow et al., 2016). Furthermore, recent studies indicate that AOA might be able to produce some oxygen internally (Kraft et al., 2022). N_2_O can be formed as a byproduct of oxygen-limited nitrification (Ji et al., 2015; Ritchie & Nicholas, 1972; Trimmer et al., 2016).

In anoxic waters and sediments, NO_3_^−^ and NO_2_^−^ can be removed via denitrification, anaerobic ammonium oxidation (anammox) or dissimilatory nitrate reduction to ammonium (DNRA). The DNRA process is carried out by phylogenetically diverse microorganisms with *nrfA* as a diagnostic gene (Cannon et al., 2019). While nitrification and DNRA retain nitrogen in a system, denitrification and anammox contribute to the loss of fixed nitrogen. Denitrification refers to the step-wise reduction of NO_3_^−^ to NO_2_^−^, nitric oxide (NO), N_2_O and dinitrogen gas (N_2_) by the proteins encoded by the *narGHI/napAB, nirK/nirS, norBC* and *nosZ* genes, respectively. These steps are often coupled to organic matter oxidation. Denitrification plays a central role in the formation as well as the removal of the greenhouse gas N_2_O, and is widespread within the domains of Bacteria and Archaea (Graf et al., 2014). In the anammox process, NH_4_^+^ and NO_2_^−^ are transformed into N_2_ via hydrazine, with hydrazine synthase (*hzsA*) as the diagnostic gene (Harhangi et al., 2012). So far three marine groups of anammox bacteria have been discovered: *Scalindua* (van de Vossenberg et al., 2008)*, Anammoxibacter* (Suarez et al., 2022), and *Bathyanammoxibiaceae* (Zhao et al., 2022). *Scalindua* appears to be more tolerant towards oxygen than freshwater species (Okabe et al., 2023).

Eutrophication and deoxygenation severely impact marine nitrogen cycling. Organic matter mineralization releases NH_4_^+^ and consumes oxygen, and subsequent oxygen depletion strongly affects the redox zonation in the sediment. Upon prolonged hypoxia, the loss of oxic and oxidized zones from the sediment will lead to an imbalance between electron acceptors, such as oxygen, NO_3_^−^ and NO_2_^−^, and electron donors, and result in the accumulation and release of NH_4_^+^ (Jäntti & Hietanen, 2012; Żygadłowska et al., 2023). Yet, the effects of eutrophication and deoxygenation on microbial nitrogen cycling in marine sediments are complex. While microbial gene abundance (Angell et al., 2018; Lipsewers et al., 2016; Wang et al., 2022), (potential) rates and processes (Jäntti & Hietanen, 2012; Peng et al., 2021; Song et al., 2021) and *in situ* geochemical profiles and fluxes (Lequy et al., 2022; Sommer et al., 2016) related to nitrogen cycling have been studied separately in coastal ecosystems impacted by eutrophication and deoxygenation, these three aspects are not often studied together nor linked.

Here, we explored the nitrogen cycling potential of sediment from a seasonally euxinic coastal ecosystem. This study aimed to (i) identify whether the nitrogen cycling potential of the sediment differs with depth and/or season (i.e. with contrasting redox conditions), (ii) explore which nitrogen cycling pathways play a role in nitrogen conversions in the sediment, and (iii) elucidate which microorganisms potentially are involved. The combined results from incubation experiments, 16S rRNA amplicon and shotgun metagenome sequencing and geochemical profiling revealed potential for nitrification, denitrification and DNRA but not anammox across depths and seasons in the sediments from this seasonally euxinic, coastal ecosystem.

## Experimental Procedures

### Sampling site and sediment collection

Lake Grevelingen is a eutrophic, coastal marine lake located in the South West of the Netherlands (Żygadłowska et al., 2023). Detailed descriptions of the lake can be found in Hagens et al., 2015, Lipsewers et al., 2016 and Żygadłowska et al., 2023. The lake has a surface area of 115 km^2^ and average water depth of 5.1 m. The lake contains several deeper basins with a maximum water depth of up to 45 m in the Scharendijke basin (Żygadłowska et al., 2023). The sedimentation rate in the basin is ∼ 13 cm year^−1^ (Egger et al., 2016). The water column in Scharendijke basin is seasonally stratified allowing the deeper water to become euxinic in summer, while it remains well-mixed and oxygenated in winter (Egger et al., 2016; Żygadłowska et al., 2023). There is a distinct seasonality in the redox chemistry of the surface sediment linked to the variations in bottom water oxygen (Klomp et al., 2025; Van Helmond et al., 2025; Żygadłowska et al., 2023). While NH_4_^+^ concentrations exceed 10 mM in the first 30 cm of the sediment in March and September, oxygen and NO_3_^−^ are depleted within a few mm in March and are not detected in euxinic sediments from September. Such seasonality is also observed for hydrogen sulfide (H_2_S), which reaches concentrations of 4 mM in the first 10 cm of the sediment in September, whereas it is detected deeper in the sediment in March.

Sediment cores were taken with a UWITEC gravity corer with transparent PVC core liners (length: 120 cm; inner diameter: 6 cm) at the Scharendijke basin (51.742 °N, 3.849 °E) during cruises with RV Navicula in March and September 2023 and March 2024. The top of the sediment was separated into 0-2 and 3-5 cm intervals and the remainder of the sediment core was sliced into 5 cm sections under N_2_ atmosphere in an anaerobic glove bag at ambient temperature. The sediment sections were stored at 4 °C in aluminum bags under N_2_ atmosphere until they were used for incubations. Sediment cores for DNA extraction were collected in March and September 2023, and samples were taken every cm up to 20 cm depth and every other cm between 20 and 40 cm depth. They were stored directly at −80 °C on RV Navicula and at −20 °C in the lab. Additional sediment cores for porosity determination were collected in March 2023 and March 2024 and sliced at a 1 cm and 0.5 cm resolution, respectively.

### Potential rates of nitrogen cycling processes

Batch incubations were set up with sediment from a range of depths to study the nitrogen cycling potential under contrasting redox conditions. In an anaerobic hood containing an N_2_:H_2_ (95/5) atmosphere, sediment was mixed 1:100 with sterile artificial seawater (ASW; 26 g/L NaCl, 5 g/L MgCl_2_ · 6 H_2_O, 1.4 g/L CaCl_2_ · 2 H_2_O, 0.5 g/L KCl, 20 mM HEPES buffer; pH set at 7.5) in 120 mL serum bottles to study the potential for nitrogen cycling processes and potential rates.

#### Aerobic nitrification potential

Aerobic nitrogen cycle processes were studied in incubations with sediment from March 2023 (0-2, 10-15 and 25-30 cm depth) and September 2023 (0-2 cm depth). The incubation bottles were left open outside the anaerobic hood and exposed to ambient air for a few minutes and were then capped. The presence of oxygen was verified by monitoring with a needle-type oxygen microsensor (PreSens, Regensburg, Germany). To study the nitrification potential of the sediment, a no substrate control (treatment 1) was compared to bottles amended with (2) 0.5 mM NH_4_Cl and (3) 0.25 mM NaNO_2_. A fourth treatment with 0.5 mM NH_4_Cl + 2% CH_4_ was set up for certain sediment sections to assess the effect of CH_4_ on the NH_4_^+^ oxidation potential, as eutrophic sediments, including those of Lake Grevelingen, are characterized by high porewater concentrations of the greenhouse gas CH_4_ (Egger et al., 2016; Żygadłowska et al., 2023, 2024). New CH_4_ (up to 5.6% of the headspace) was added to these bottles upon CH_4_ depletion. Because the oxic incubations with sediment from 0-2 cm depth from March 2023 may have suffered from oxygen depletion, these incubations were repeated with sediment from 0-2 cm depth from March 2024. Moreover, a fifth treatment was included for this sediment section, comprising a 1:100 dilution of the sediment in cotton-plugged Erlenmeyers supplemented with 0.5 mM NH_4_Cl. The bottles were incubated for 4 weeks or until substrate depletion.

#### Fate of NO_2_^−^ under oxic conditions

Because the concentration of NO_3_^−^ formed upon the oxidation of NH_4_^+^ and NO_2_^−^ was lower than expected based on the stoichiometry of NH_4_^+^ and NO_2_^−^ oxidation (Figure S1), additional oxic incubations were prepared in March 2024 to assess the fate of NO_2_^−^ in these bottles. Sediment from 25-30 cm depth was selected as it had shown good NO_2_^−^ oxidation potential (Figure S1). The bottles were amended with 0.2 mM Na^15^NO_2_ (treatment 6), and they were incubated for 4 weeks.

#### Potential for anaerobic nitrogen cycle processes

Anaerobic nitrogen cycle processes were studied in incubations with sediment from March 2023 (0-2, 10-15 and 25-30 cm depth) and September 2023 (0-2 cm depth). The incubation bottles were capped and their headspace was exchanged with argon gas with overpressure by subsequent rounds of gassing and vacuum after taking the bottles out of the anaerobic hood. The absence of oxygen was verified with a needle-type oxygen microsensor (PreSens, Regensburg, Germany). Again, a no-substrate control (treatment 7) was compared to bottles amended with (8) 1 mM NaNO_3_, (9) 0.25 mM NaNO_2_ + 0.5 mM NH_4_Cl + 2% CO_2_, and (10) 0.5% N_2_O. Additional treatments (11) 1 mM NaNO_3_ + 0.5 mM monomethylamine (CH_3_NH_2_, MMA), and (12) 0.5% N_2_O + 0.5 mM MMA were included to study the effect of these carbon compounds on NO_3_^−^ and N_2_O reduction. MMA is a common metabolite in marine systems and forms a direct link between the carbon and nitrogen cycles (Mausz & Chen, 2019). Because 1 mM NO_3_^−^ was not fully converted and resulted in the accumulation of NO_2_^−^ in the treatment 8 incubations made with sediment from March 0-2 cm (Figure S2B), 0.5 mM NaNO_3_ was added in treatment 8 and 11 for the incubations with the other sediment layers. The bottles were incubated for 4 weeks or until substrate depletion.

#### Anammox potential

To check for anammox potential in the sediment, incubations were set up with sediment from the 25-30 cm section from March 2024, which was characterized by high NH_4_^+^ and low sulfide concentrations (Żygadłowska et al., 2023). Incubation bottles were amended with (13) 0.2 mM Na^14^NO_2_ + 0.2 mM ^15^NH_4_Cl and (14) 0.2 mM Na^15^NO_2_. Similar incubations were set up with *Scalindua* biomass taken from an in-house reactor to verify if anammox could be detected with this set-up (Russ et al., 2014; van de Vossenberg et al., 2008). The marine *Scalindua* cells were washed with ASW and centrifuged at 2000 x g for 10 minutes at room temperature, until no more NO_2_^−^ and NO_3_^−^ were detected. The cells were diluted 10x in artificial sea water in 120 mL serum bottles and amended with (15) 0.2 mM Na^14^NO_2_ + 0.2 mM ^15^NH_4_Cl (16) 0.2 mM Na^15^NO_2_ + 0.2 mM ^14^NH_4_Cl. In treatment 17, the 10x diluted *Scalindua*-containing biomass was mixed with 1:100 diluted sediment from 25-30 cm depth from March 2024 and amended with only 0.2 mM Na^15^NO_2_. Because of the high background NH_4_^+^ levels in these bottles, no supplementary ^14^NH_4_Cl was added to the treatment 14 and 17 bottles. The bottles were incubated for 48 hrs.

#### Relevance of abiotic NO_2_^−^ removal

To assess the role of abiotic conversions in NO_2_^−^ removal in the sediments, bottles with sediment from 25-30 cm depth from March 2024 were autoclaved twice and subsequently amended with 0.2 mM Na^15^NO_2_ in oxic and anoxic conditions (treatments 18 and 19, respectively). The bottles were incubated for 48 hrs and 28 days, respectively.

An overview of all treatments is given in Table S1. For all treatments, biological duplicates were set up. Because of the high reproducibility of the background processes occurring in the no substrate controls in the March 0-2 and 25-30 cm incubations (Figure S1, Figure S2), biological duplicates were omitted for the no substrate controls (treatment 1 and 7) in the March 10-15 cm and September 0-2 cm incubations. Likewise, treatment 5 was performed with one bottle only with sediment from March 10-15 cm and September 0-2 cm. All bottles were incubated in the dark at 21 °C on a shaker at 100 rpm.

### Chemical analyses

#### Nitrogen compounds

Liquid samples for NO_2_^−^, NO_3_^−^, NH_4_^+^ and MMA determination were taken from the incubations over time. The samples were centrifuged for 5 minutes at 20238 x g and the supernatant was stored at −20 °C until analyzed. NO_2_^−^ and NO_3_^−^ levels were monitored using colorimetric test strips for NO_2_^−^ and NO_3_^−^ (MQuant®) and quantified colorimetrically in triplicate by the Griess assay (Miranda et al., 2001). NH_4_^+^ concentrations were measured in triplicate by fluorescence upon reaction with the *o*-phtaldehyde (OPA) reagent containing 4 mM OPA and 0.7 mM 2-mercaptoethanol buffered with 400 mM potassium phosphate buffer at pH 7.3-7.4 (Goyal et al., 1988). The reagent was mixed in a 15:1 ratio with the samples. Fluorescence was measured by a Spark M10 plate reader (Tecan, Männedorf, Switzerland) at an excitation wavelength of 411 nm and emission wavelength of 482 nm.

#### MMA determination

To measure MMA, a derivative was formed with benzaldehyde. To this end, 0.4% benzaldehyde in 70% acetonitrile and 83 µM NaOH, sample and Milli-Q water were mixed in a 2:1:1 ratio and this mixture was incubated for 1 hr at room temperature in the dark after vortexing for 10 seconds. The products were extracted from the samples with dichloromethane in a 1:1 ratio, and measured by a 7890A gas chromatograph (GC; Agilent Technologies, Santa Clara, CA, USA) containing an HP-5MS column (30 m x 0.25 mm x 0.25 μm) and equipped with an 7693A autosampler. The GC was coupled to a JEOL JMS-T100 GCv mass spectrometer (JEOL Ltd., Akishima, Tokyo, Japan).

#### Gas and GC-MS determination

Headspace gas samples were taken by a gastight Hamilton syringe. Headspace N_2_O concentrations were measured over time by an HP6890 Plus GC (Agilent, Santa Clara, CA, USA) containing an HP 6 FT Porapak Q 60/100 column and equipped with an electron capture detector. Headspace concentrations of labeled gases were measured over time with an 8890 GC (Agilent, Santa Clara, CA, USA) containing an Agilent 6 FT Porapak Q 80/10 column and coupled to a 5977B mass selective detector (Agilent, Santa Clara, CA, USA).

### Calculations

#### Dissolved gases

To calculate the amount of dissolved N_2_O and N_2_, the Bunsen coefficient (β) was calculated by (eq. 1), taking into account the salinity (S) and temperature (T) (Weiss, 1970; Weiss & Price, 1980). The specific Bunsen coefficients A_1-3_ and B_1-3_ for N_2_O and N_2_ were taken from Weiss & Price (1980) and Weiss (1970), respectively. The Henry solubility coefficient (H^cp^) was calculated by (eq. 2), with R the ideal gas constant, and T^STP^ the standard temperature for the Bunsen coefficient (273.15 K) (Sander, 2015). The concentration of dissolved gas (c_a_) was calculated by multiplying the Henry solubility constant with the partial pressure (p_g_) of the gas of interest (eq. 3).

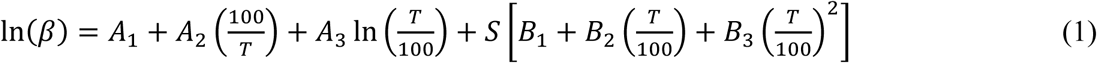

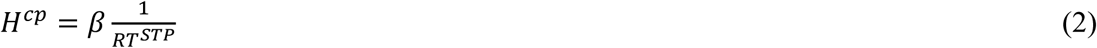

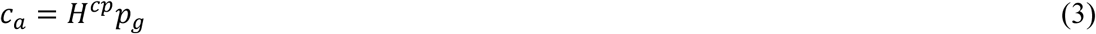

The maximum potential rates of reduction and/or oxidation of NH_4_^+^, NO_2_^−^, NO_3_^−^ and N_2_O (r_x_) were calculated by eq. 4, where c is the concentration of the compound of interest at t_i_ or t_j_, t_i_ and t_j_ are the timepoints between which the slope in the plots of time vs c is steepest, V is the liquid volume in the incubation bottle at the moment of sampling at t_i_ or t_j_, and m is the amount of sediment used to inoculate the bottle converted to dry weight. Based on the porosity samples, the percentage dry weight of the sediment was 17.7%, 24.1%, 25.4% for March 2023 0-2, 10-15, 25-30 cm and 22.8% for March 2024 0-2 cm, respectively. For the September incubations, the porosity from the surface sediments from March 2023 was used. 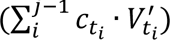 accounts for the amount of the compound of interest taken out by liquid samples between t_i_ and t_j_, with V’t_i_ the volume of liquid sample taken at timepoint t_i_. This was considered neglectable for N_2_O and was therefore not taken into account in the calculation of the N_2_O reduction rates. Because NO_2_^−^ was also removed anaerobically in the oxic bottles, the potential rates of NO_2_^−^ oxidation were calculated based on the amount of NO_3_^−^ formed.

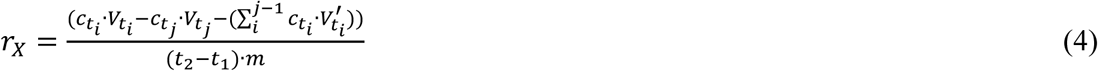

#### Mass balance

To determine which processes contribute to NO_2_^−^ removal in oxic conditions, a mass balance was set up according to eq. 5 and 6. The total amount of N in moles at the start and end of the incubations (N_i_ and N_f_, respectively) was calculated by adding up the total amount of moles of N present in NH_4_^+^, NO_2_^−^, NO_3_^−^, and all isotopes of N_2_O and N_2_ at both timepoints, corrected for the amounts N taken out by taking gas and liquid samples.

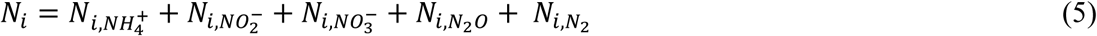

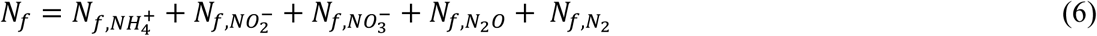

#### Contribution of DNRA and denitrification

No net background production of NH_4_^+^ was observed in the anoxic no-substrate treatment for all four sediment sections (Figure S2), and anammox did not contribute to NO_3_^−^ and NO_2_^−^ removal in the sediment incubations (Figure S3). Therefore, the relative contribution of DNRA (R_DNRA_) and denitrification (R_dnf_) to the observed NO_3_^−^ and NO_2_^−^ removal was estimated for each sediment section from treatments 8 and 9 by eq. 7 and 8, where [NO_x_^−^]_begin_ is the NO_3_^−^ or NO_2_^−^ concentration at the start of the incubations. In the presence of MMA, the contribution of DNRA relative contribution of DNRA (R_DNRA,MMA_) and denitrification (R_dnf,MMA_) to the observed NO_3_^−^ removal was calculated by eq. 9 and 10, where the difference in the concentration of MMA at the start and end accounts for the amount of NH_4_^+^ produced upon MMA oxidation.

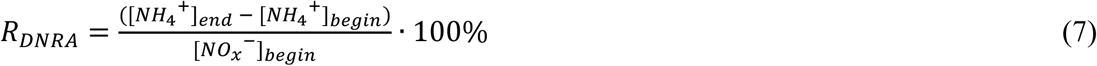

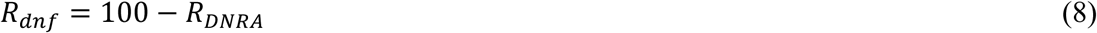

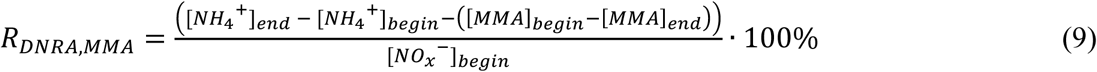

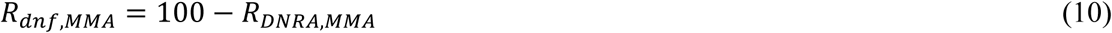

### DNA extraction

Samples for DNA were taken from the treatment 1-5 and 7-12 bottles at the end of incubation. Upon centrifugation for 2 minutes at 10000 x g, the pellet was stored at −20 °C until DNA extraction. DNA was extracted with the DNeasy PowerSoil Pro Kit (Qiagen, Venlo, the Netherlands) following the manufacturer’s instructions. The cells were lyzed by bead beating in a tissue disruptor (Qiagen, Venlo, the Netherlands) at 50 Hz for 10 minutes.

### 16S rRNA amplicon sequencing and data analysis

Extracted DNA was sent for sequencing of the V3-V4 16S rRNA region (Illumina MiSeq platform, Macrogen, Seoul, South Korea) with the primers Bac341F (CCTACGGGNGGCWGCAG; Herlemann et al., 2011) and Bac806R (GGACTACHVGGGTWTCTAAT; Caporaso et al., 2012) for bacteria and Arch349F (5′-GYGCASCAGKCGMGAAW-3′) and Arch806R (5′-GGACTACVSGGGTATCTAAT-3′) for archaea (Takai & Horikoshi, 2000). The raw reads were processed according to the DADA2 pipeline (Callahan et al., 2016) and the microbial community was analyzed by the phyloseq package (v1.44.0; McMurdie & Holmes, 2013) in RStudio (R v4.3.1) as described by Venetz et al. (2023). The bacterial reads were trimmed to a length of 270 and 215 nt for forward and reverse reads, respectively. The archaeal forward and reverse reads were trimmed to lengths of 300 and 200 nt, respectively. The taxonomy was assigned based on the Silva database (v138.1; Quast et al., 2013). For the bacterial reads, only ASVs classified as Bacteria at the Kingdom level, and not as the Order Chloroplast or Family Mitochondria were used for downstream processes. Likewise, solely the ASVs classified as the Kingdom Archaea were selected in the analysis of the archaeal sequences. The results were visualized by ggplot2 (v.3.5.0; Wickham, 2016). The 16S rRNA raw sequencing reads are deposited in NCBI under BioProject number PRJNA1215350.

### Metagenome sequencing and data analysis

To identify the genomic potential for different nitrogen cycling processes in the studied sediment sections, DNA extracted from these sediment sections was sent for PCR-free metagenome sequencing (Illumina DNA PCR-free library preparation (450 bp insert size), NovaSeq X platform, Macrogen, Amsterdam, the Netherlands). SingleM (Woodcroft et al., 2024) was run on the raw reads to identify the percentage archaea that made up the total microbial community and to analyze the archaeal community composition of the sediment. Sequencing adapters were removed from the reads, alongside quality trimming (at Q15) and length filtering (minimum length 100 bp) with BBDuk (v37.76; BBTools package, Bushnell, 2014). The reads were then error-corrected using Tadpole (k=50; BBTools) and normalized with BBNorm (target=30, min=2; BBTools). Assembly was done for each sample separately with Metaspades (v3.15.5; Nurk et al., 2017), and the reads were mapped to the assembly by BBMap (slow=t; BBTools). Annotation of the contigs and subsequent analysis of the nitrogen cycle genes was done with Metascan using the --nokegg flag (Cremers et al., 2022). Metascan was also run with the HMM profile of DNA gyrase subunit A (*gyrA*; COG0085 (bacteria) from EggNog 5.0.0), as a single-copy marker gene for the estimation of the total abundance of cells. The taxonomy of the identified *amoA*, *nxrA* and *nosZ* genes was determined by a BLASTP search of their protein sequences against the nr database. The coverage of the contigs in reads per million (RPM) was calculated by coverM (Aroney et al., 2025), and per sample the number of occurrences and the coverage per occurrence were summed for each gene. The results were visualized by ggplot2 (v.3.5.0; Wickham, 2016). The raw metagenome sequencing reads are deposited in NCBI under BioProject number PRJNA1215350.

## Results

### Potential nitrification rates

The microbial nitrogen cycling potential in sediment from Lake Grevelingen was explored in batch incubations with sediment from different depths and seasons, to capture contrasting redox conditions. Surprisingly, potential for NH_4_^+^ oxidation was observed in all four sediment sections, including those that are anoxic *in situ*. Sequential formation of NO_2_^−^ and NO_3_^−^ was observed upon oxidation of both endogenous and supplemented NH_4_^+^ in incubations exposed to air (Figure 1; Figure S1). After a lag phase of 7-10 days, the maximum potential rates of NH_4_^+^ oxidation ranged from 16 to 21 µmol g^−1^ day^−1^ across depths in March, and were highest in the top sediment from September (53 µmol g^−1^ day^−1^; Figure 2). The addition of CH_4_ reduced the maximum potential rate of NH_4_^+^ oxidation, particularly in surface sediment from September where it was reduced to 4.8 µmol g^−1^ day^−1^ (Figure S4). CH_4_ was also oxidized in these bottles. Maximum potential NO_2_^−^ oxidation rates ranged from 2.0-2.8 µmol g^−1^ day^−1^ in incubations with anoxic sediments to 21 µmol g^−1^ day^−1^ in the top sediment in March (Figure 2).

**Figure 1.**
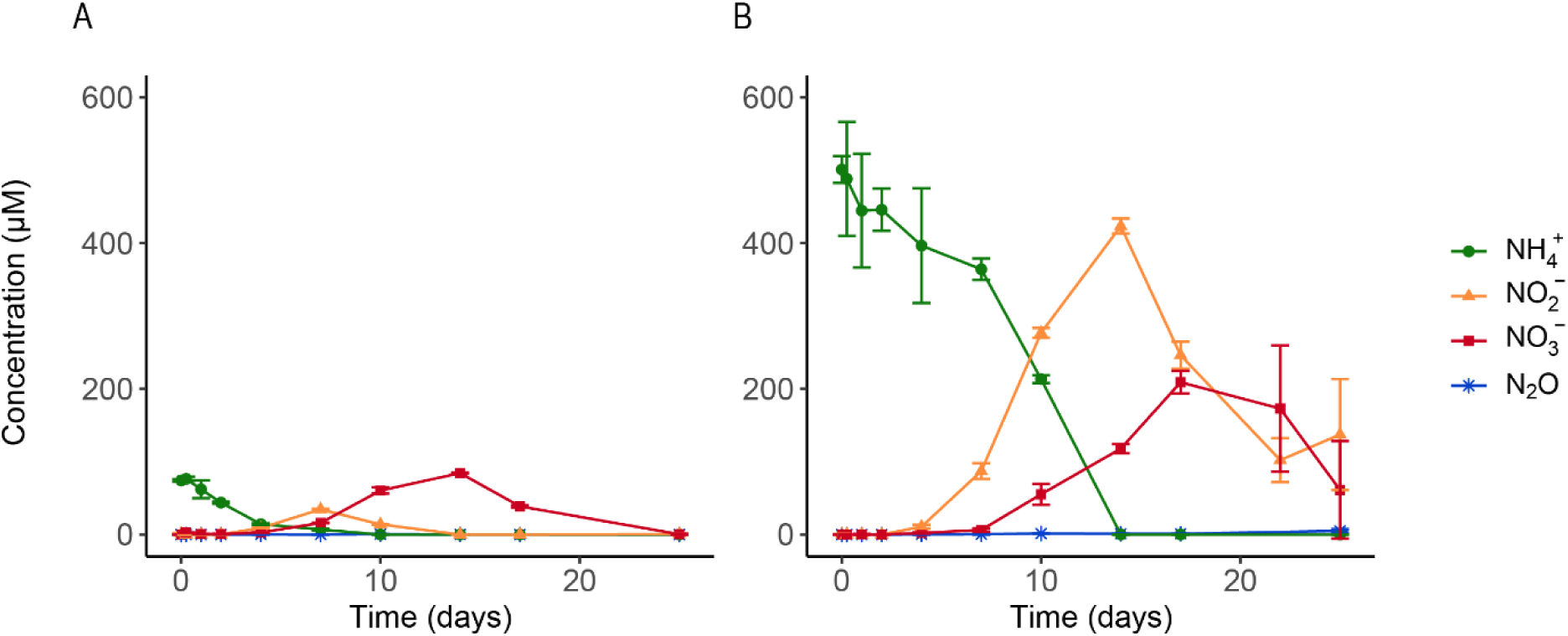
Concentrations of dissolved NH_4_^+^ (green), NO_2_^−^ (orange), NO_3_^−^ (red), and headspace N_2_O (blue) over time in bottle incubations with sediment from Lake Grevelingen from March 10-15 cm depth supplemented with (A) no substrate (treatment 1) and (B) 0.5 mM NH_4_Cl (treatment 2) after exposure to ambient air. Error bars indicate the standard deviation of the triplicate measurements for each sediment-treatment combination.

**Figure 2.**
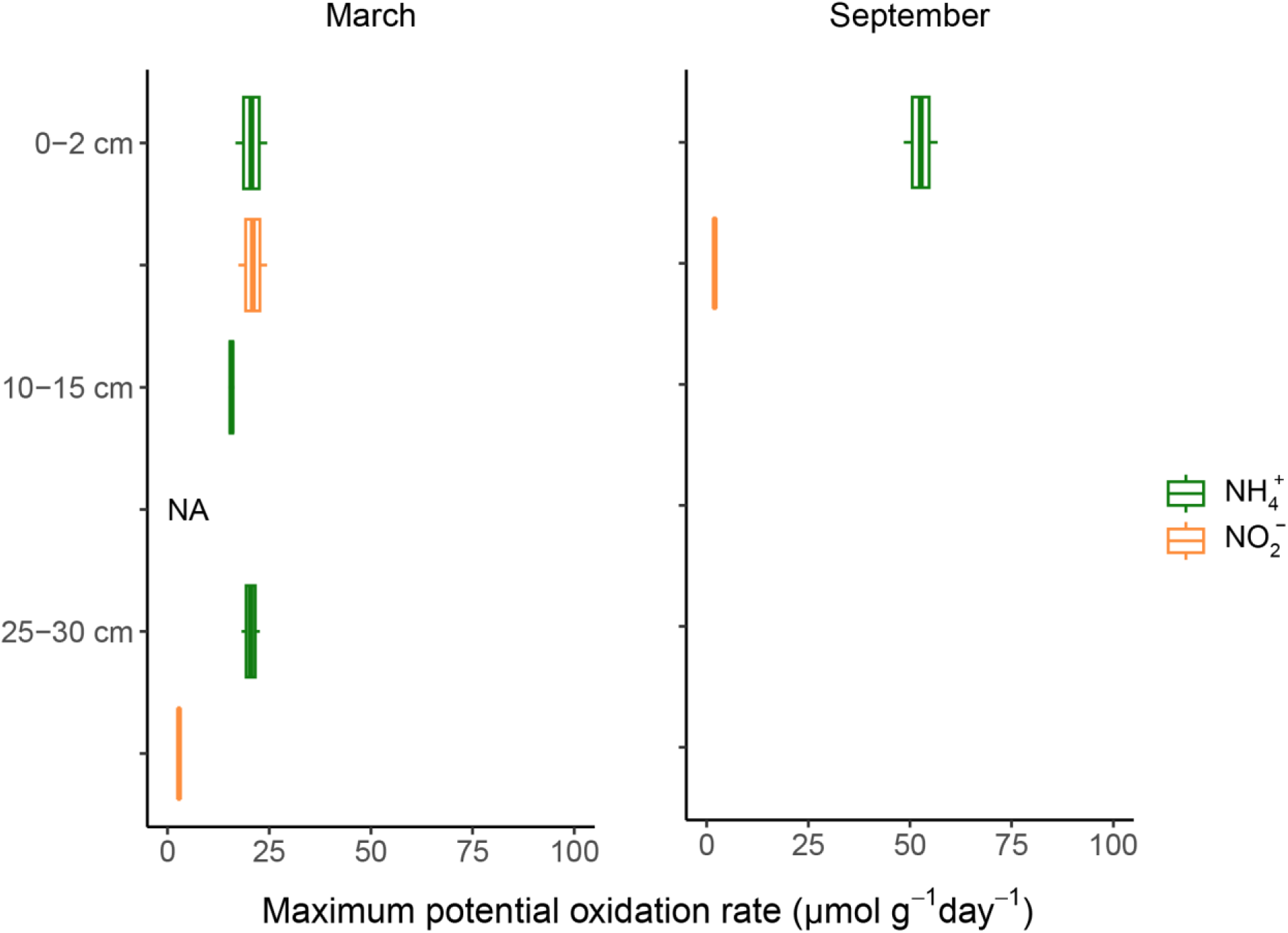
Maximum potential rates (µmol g^−1^ day^−1^) of NH_4_ ^+^ (green) and NO_2_ ^−^ (orange) oxidation at different sediment depths in March and September, as measured in duplicate bottles amended with 0.5 mM NH_4_Cl and 0.25 mM NaNO_2_ (treatments 2 and 3, respectively). The maximum potential rate of NO_2_ ^−^ oxidation was calculated based on the formation of NO_3_ ^−^. NA: treatment not included for incubations with this sediment section.

### Processes contributing to NO_2_^−^ loss under oxic conditions

Upon the oxidation of 0.5 mM NH_4_^+^ and the resulting NO_2_^−^, less than the expected 0.5 mM NO_3_^−^ was formed in most sediment sections (Figure S1B). To assess the processes contributing to NO_2_^−^ loss under oxic conditions, separate oxic incubations supplemented with ^15^NO_2_^−^ were performed (treatment 6). Again, a disparity in N compounds was observed, as 20.4-24.1 µmol N was present in NH_4_^+^ and NO_2_^−^ at the start of the incubations, while only 7.2-8.3 µmol N was retrieved in NO_2_^−^ and NO_3_^−^ at the end of the incubations (Table S2). Formation of some ^45^N_2_O and ^46^N_2_O was observed over time, as well as a slight increase (0.02-0.10 µmol N) in ^30^N_2_ at the end of the incubations (Table S2, Figure S5). Autoclaved controls showed no removal of NO_2_^−^ (Figure S6A).

### Potential rates of NO_3_^−^, NO_2_^−^ and N_2_O reduction

All four studied sediment sections showed high potential for NO_3_^−^, NO_2_^−^ and N_2_O reduction under anoxic conditions (Figure 3, Figure S2). Upon NO_3_^−^ reduction, both formation and removal of NO_2_^−^ and N_2_O were observed. Moreover, NH_4_^+^ was produced over time in treatments 8 and 9 upon reduction of NO_3_^−^ and NO_2_^−^ (Figure S2B, C). Maximum potential NO_3_^−^ reduction rates ranged from 55 to 84 µmol g^−1^ day^−1^ in March and peaked at 167 µmol g^−1^ day^−1^ in the surface sediment of September (Figure 4). Reduction rates decreased from oxidized to more reduced species in all studied sediment sections, except for 0-2 cm depth in March, where the maximum potential rate of NO_2_^−^ reduction exceeded that of NO_3_^−^ reduction. Supplementation of MMA as a carbon and energy source did not change the potential rates for NO_3_^−^ and N_2_O reduction (Figure S7), indicating sufficient endogenous electron donor. Yet, the addition of MMA stimulated full conversion of 1 mM NO_3_^−^ in the 0-2 cm March incubations (Figure S2E).

**Figure 3.**
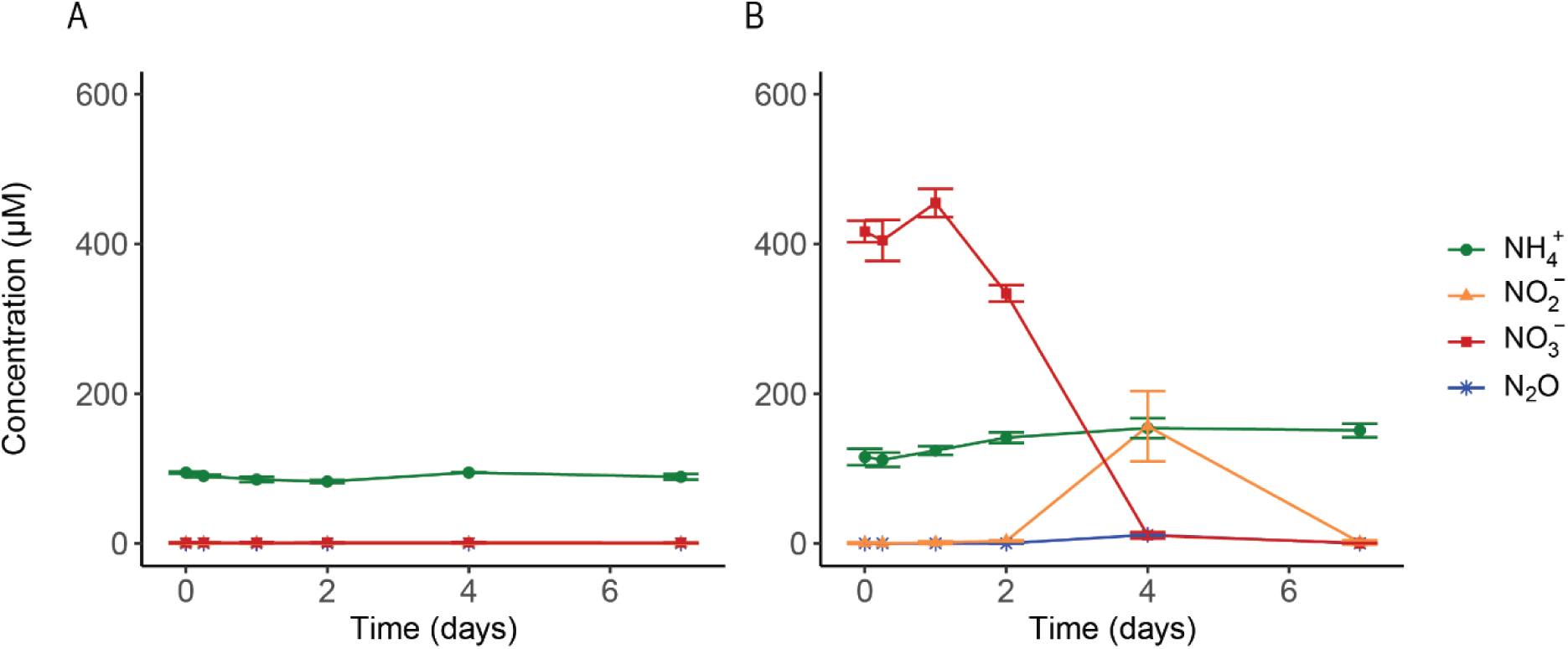
Concentrations of dissolved NH_4_^+^ (green), NO_2_^−^ (orange), NO_3_^−^ (red), and headspace N_2_O (blue) over time in incubations with sediment from Lake Grevelingen from March from 10-15 cm depth supplemented with (A) no substrate (treatment 7) and (B) 0.5 mM NaNO_3_ (treatment 8). Error bars indicate the standard deviation of the triplicate measurements for each sediment-treatment combination.

**Figure 4.**
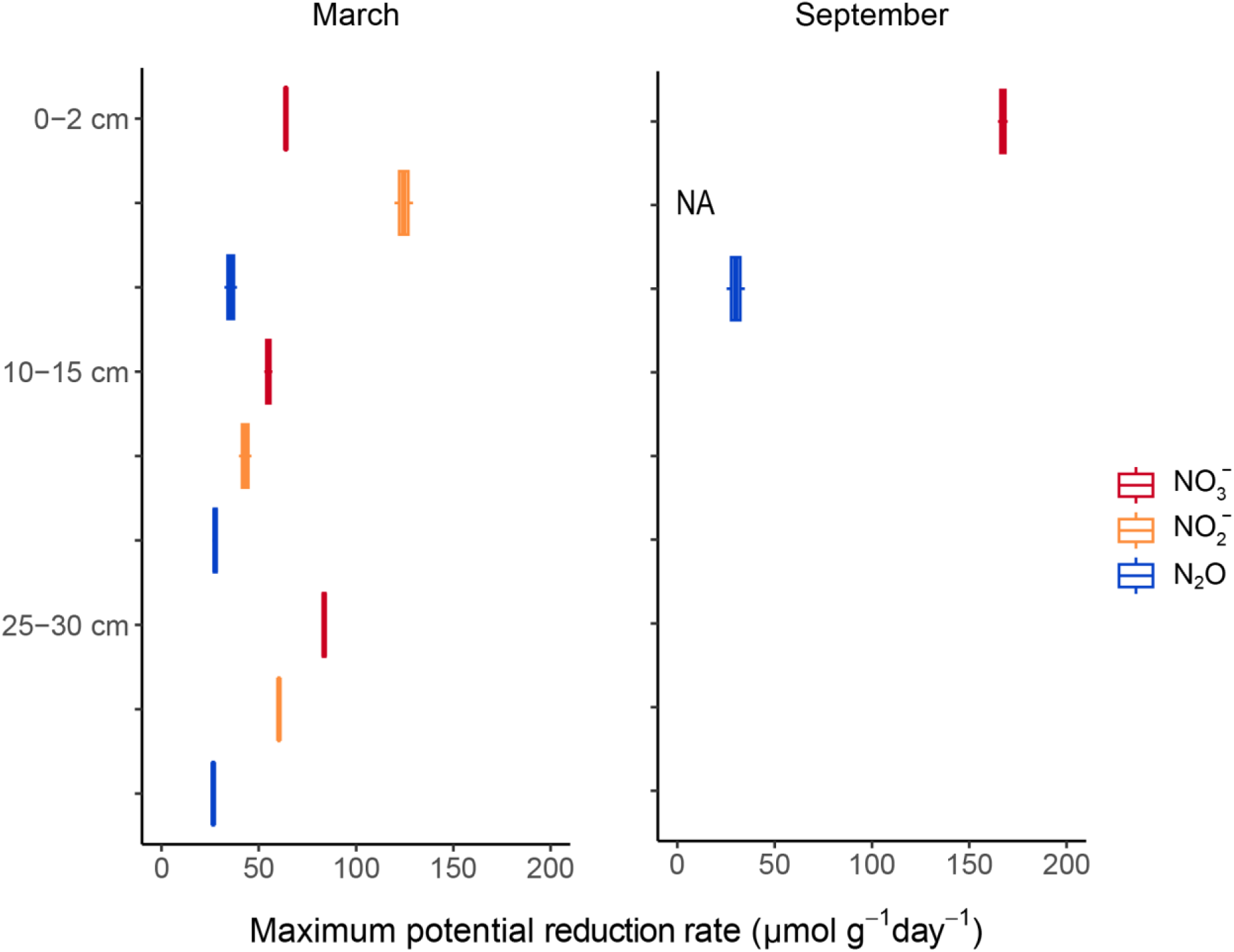
Maximum potential rates (µmol g^−1^ day^−1^) of NO_3_ ^−^ (red), NO_2_ ^−^ (orange) and N_2_O (blue) reduction at different sediment depths in March and September, as measured in duplicate incubation bottles amended with 0.5-1 mM NaNO_3_ (treatment 8), 0.25 mM NaNO_2_ + 0.5 mM NH_4_Cl + 2% CO_2_ (treatment 9), and 0.5% N_2_O (treatment 10), respectively. NA: treatment not included for incubations with this sediment section.

### Contribution of DNRA, denitrification and anammox to anaerobic NO_3_^−^ reduction

No measurable anammox rates were observed in incubations with sediment (Figure S3). Formation of ^29^N_2_ was detected in incubations with and without sediment inoculated with external supplemented biomass of *Scalindua*, indicating that the experimental set-up was suitable to detect anammox activity and that the slurries did not contain inhibitory compounds for anammox. Thus, we consider that NO_3_^−^ reduction proceeded via DNRA and denitrification. In all studied sediment sections, denitrification contributed more to NO_3_^−^ reduction than DNRA. The relative contribution of DNRA was 5.6% and 1.6% in the surface sediments of March and September, respectively, and increased with depth to 20.7% (Figure 5). The relative contribution of DNRA was higher for the reduction of NO_2_^−^ in the presence of 0.5 mM NH_4_^+^ for all sediment depths in March, and peaked at 38.9 % at 25-30 cm depth (Figure S8A). Upon MMA addition the relative contribution of DNRA increased to 11.4 and 38.2% for the 0-2 and 25-30 cm sediment sections from March, respectively (Figure S8B). In anoxic, abiotic controls supplemented with NaNO_2_, no removal of NO_2_^−^ was observed (Figure S6B).

**Figure 5.**
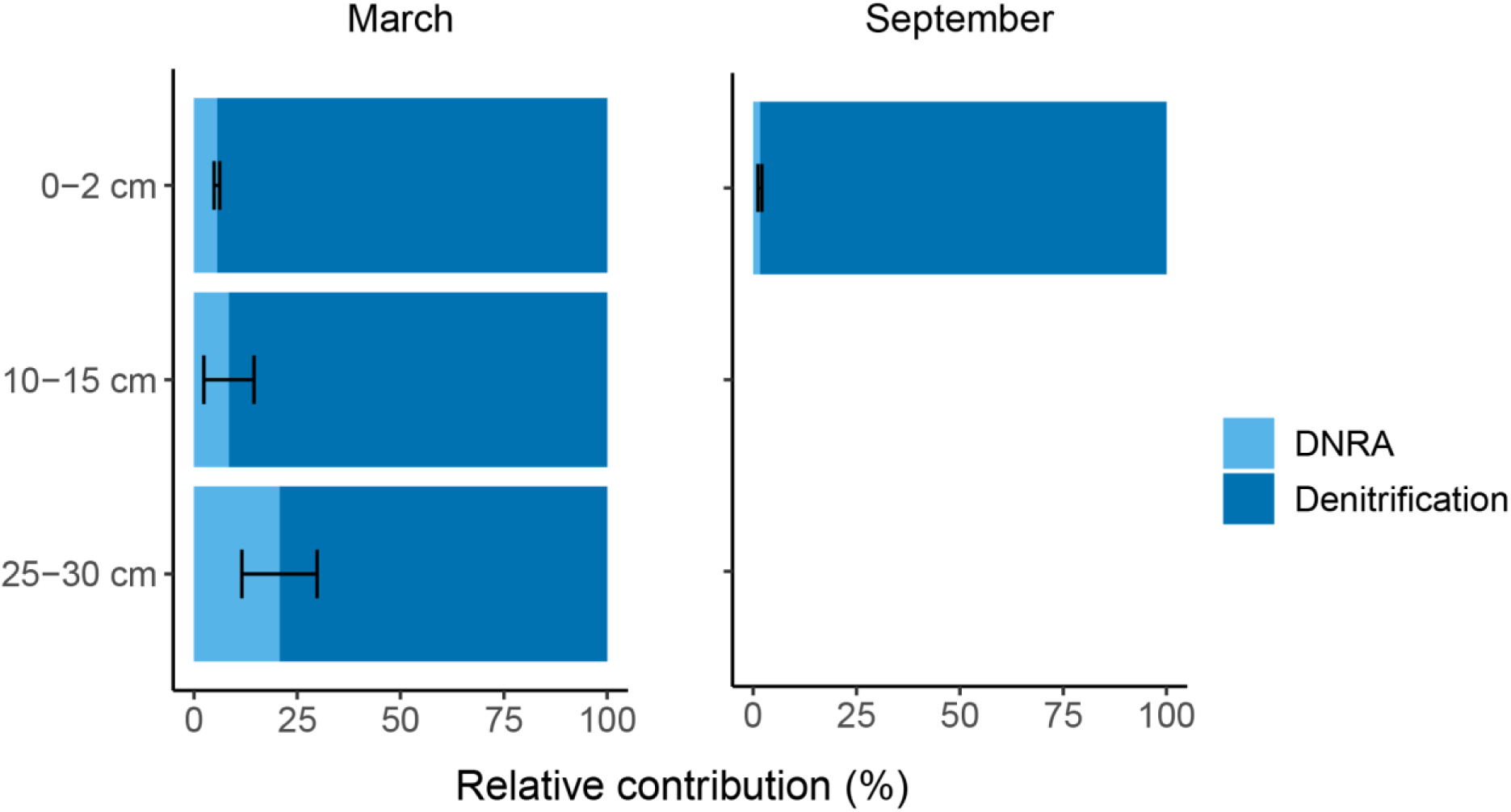
Relative contribution of DNRA and denitrification to NO_3_ ^−^ reduction at different sediment depths in March and September as measured in bottles amended with 0.5-1 mM NaNO_3_ (treatment 8). Error bars indicate the standard deviation between duplicate bottles.

### Composition of the nitrogen cycling microbial community

The microbial community composition of the initial sediment as well as that after incubation was characterized by 16S rRNA amplicon and shotgun metagenome sequencing to identify which microorganisms were present in each sediment section and enriched under the specific applied condition. All four sediment sections were dominated by the bacterial phyla Proteobacteria (22.5-35.0%), Desulfobacterota (10.4-21.0%) and Bacteroidota (11.2-17.9%) (Figure S9, Figure S10). The sediment was highly diverse with 16.7-29.8% of the reads mapped to the rare biosphere ‘others’ (< 2%) at the family level (Figure S11, Figure S12). The most dominant family in all sediment sections was *Flavobacteriaceae* (7.9-16.2%). The relative abundance of the total bacterial nitrifying community was higher in deeper sediments (0.99-1.4%) than in surface sediments (0.19-0.42%), and was dominated by *Nitrospiraceae*, *Nitrosomonadaceae* and *Nitrosococcaceae* (Figure S13). Based on the metagenome sequencing data, archaea comprised 1.7-2.8% of the total microbial community in the surface sediments and 5.0-6.6% in the deeper sediments. The Nanoarchaeota were the dominant archaeal phylum in the anoxic sediments (up to 63.2% relative abundance of archaea) (Figure S14). The AOA family *Nitrosopumilaceae* was detected in the sediment sections of March up to 14.3% relative abundance of archaea, whereas its relative abundance was 0.84% in the surface sediments in September (Figure S15).

Under oxic conditions, Proteobacteria became more dominant, reaching relative abundances up to 80.3% (Figure S9). Accordingly, the addition of oxygen to the incubations lacking CH_4_ led to an increase in relative abundance of Proteobacteria families such as *Rhodobacteraceae* and *Rhizobiales Incertae sedis* in all sediment sections and *Nitrincolaceae* and *Labraceae* in incubations with sediment from March up to 27.2%, 22.5%, 39.1% and 24.0%, respectively (Figure S11). Despite the addition of different substrates to the oxic incubations, the microbial community compositions for the oxic incubations were similar for each sediment section. Upon the addition of CH_4_ (treatment 4), the methanotrophic *Methylomonadaceae* and methylotrophic *Methylophagaceae* became more abundant compared to the original sediments and the oxic bottles amended with NH_4_^+^ only, reaching relative abundances of 7.8-34.3% and 0.2-6.9%, respectively. The relative abundance of the nitrifying bacterial community in the oxic incubations with NH_4_^+^ (treatment 2) was similar or higher and slightly enriched in *Gallionellaceae* (Candidatus Nitrotoga), *Nitrosomonadaceae, Nitrospinaceae* and *Nitrospiraceae* compared to the corresponding sediment (Figure S13). While *Nitrosopumilaceae* was one of the dominant archaeal families in the oxic incubations with NH_4_^+^ (treatment 2) with surface sediments from March, methanogenic *Methanosarcinaceae, Methanomicrobiaceae* and *Methanosaetaceae* were the dominant archaeal families in these incubations inoculated with anoxic sediments (Figure S15).

The anoxic incubations supplemented with oxidized nitrogen species were characterized by an increased relative abundance of Campylobacterota up to 43.6%, particularly upon supplementation with oxidized nitrogen species (Figure S10). Accordingly, the relative abundance of the Campylobacterota families *Arcobacteraceae* and *Sulfurimonadaceae* in these incubations was also higher compared to the sediment and the no substrate control for the surface and deeper sediments in March, respectively, particularly in the bottles supplemented with N_2_O (Figure S12). The addition of NO_3_^−^ to the anoxic incubations (treatment 8) resulted in the enrichment of *Desulfuromonadaceae* with depth and *Nitrincolaceae* for all sediment sections albeit particularly in the surface sediments of September. In line with the similar NO_3_^−^ reduction rates in the presence and absence of MMA (treatments 8 and 11), similar microbial community compositions were observed for both treatments. Consistent with the absence of anammox activity, only 2 reads of *Candidatus* Brocadia were detected in the bottles with sediment from March 25-30 cm depth amended with NO_3_^−^, but not in any of the other samples.

### Distribution of nitrogen cycle genes

Metagenome sequencing was done to identify the distribution of nitrogen cycle genes in the sediment sections. The genes encoding for the complete denitrification (*narG/napA, nirK/nirS, norB, nosZ*) and DNRA (*nrfA/otr*) pathways were detected at all sediment sections, yet were less abundant than the single-copy *gyrA* marker gene (Figure 6). *nirS* was more abundant than *nirK*. Most of the detected *nosZ* genes were classified as ‘Sec-nosZ’ and were similar to those from members of the Flavobacteriia and Gammaproteobacteria. In line with the absence of anammox from the microbial communities, *hzs* was not detected. Nitrification genes were detected in the deeper sediments from March only. The detected *amoA* genes were classified as *amoA* from *Nitrosomonadaceae* and *Nitrosopumilaceae*, whereas the *nxrA* genes belonged to Nitrospirota bacteria.

**Figure 6.**
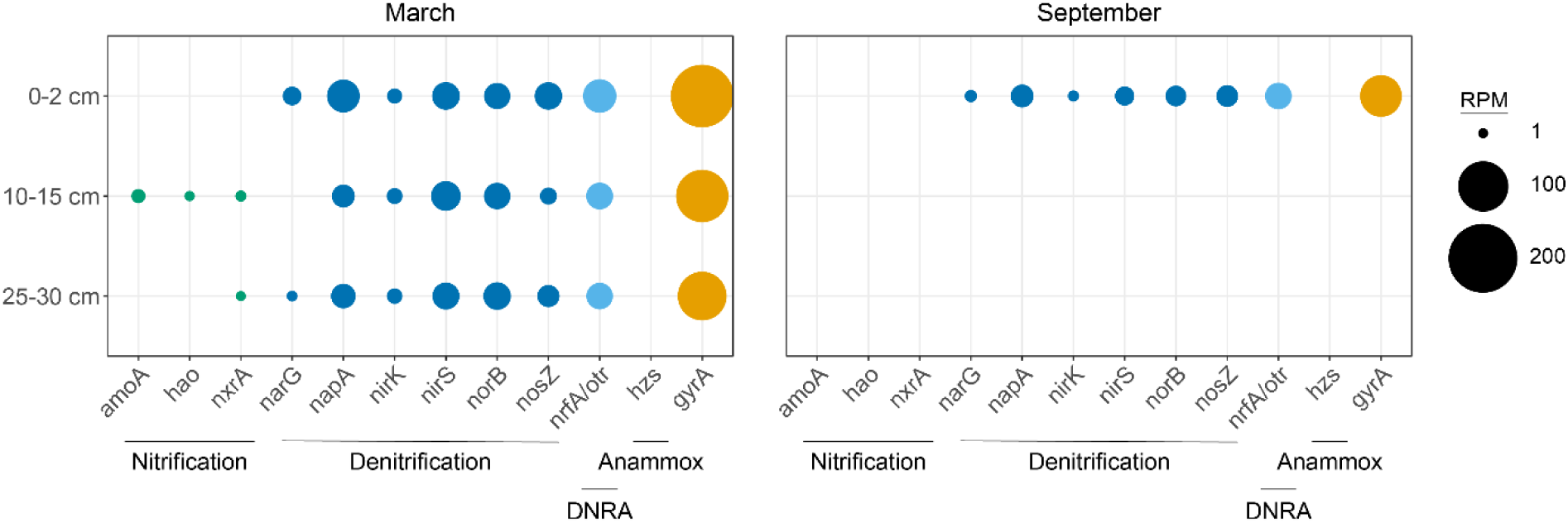
The relative abundance of genes belonging to the nitrification, denitrification, DNRA and anammox processes in reads per million (RPM) at the different depths and seasons studied as determined by metagenome sequencing. Left panel: March. Right panel: September.

## Discussion

### Potential for NH_4_^+^ and NO_2_^−^ oxidation in anoxic sediments

In this study, the nitrogen cycling potential of sediments from the seasonally euxinic Lake Grevelingen was investigated. This sediment was characterized by the rapid depletion of electron acceptors such as oxygen, NO_3_^−^ and NO_2_^−^ (NO_x_) and by accumulation of H_2_S in surface sediments in summer and deeper sediments in winter. To capture these different redox zones, sediments from various depths and two seasons (winter and summer) were used. The nitrogen cycling potential was assessed by activity measurements in batch incubations and by the relative abundance of nitrogen cycling microorganisms and their nitrogen cycling genes. High NH_4_^+^ concentrations were observed in all layers, but no specific zone of NH_4_^+^ removal was identified in the sediment. These findings are in contrast with the redox zone-specific CH_4_ filter located at 5-15 cm depth in Lake Grevelingen, where sulfate-dependent anaerobic CH_4_ oxidation occurred (Wallenius et al., 2024). Surprisingly, potential for and revival of NH_4_^+^ oxidation was observed in all four selected sediment sections at rates between 16 to 53 µmol g^−1^ day^−1^. These rates are comparable to the NH_4_^+^ oxidation rates found in 48-hr incubations for the sediments of the Colne estuary (Li et al., 2015) but considerably higher than the NH_4_^+^ oxidation rates measured in 24-hr incubations in sediments from two contrasting coastal bays in Ireland (Duff et al., 2017; Zhang et al., 2018). It should be noted that the maximum potential rates measured in this study were measured after a lag phase of 7-10 days for most sediment sections, preceded by NH_4_^+^ oxidation at much lower rates. Hence, the rates presented here are likely to be an overestimation of the *in situ* NH_4_^+^ oxidation rates.

In line with the NH_4_^+^ oxidation rates across the studied depths and seasons, nitrifying microorganisms were detected in all sediment sections. *Nitrospiraceae* were detected in all sediment sections but the resolution of the 16S rRNA sequencing data is not high enough to determine whether these are comammox *Nitrospira*. The addition of CH_4_ to the oxic incubations amended with NH_4_^+^ resulted in (slightly) lower NH_4_^+^ oxidation rates and an increase in the relative abundance of methanotrophs from the *Methylomonadaceae* family and methylotrophs belonging to the *Methylophagaceae* family. CH_4_ can also occupy the ammonium monooxygenase (AMO) catalytic pocket and can thus inhibit NH_4_^+^ oxidation (Oudova-Rivera et al., 2023; Suzuki et al., 1976; Ward, 1990).

NO_2_^−^ accumulated upon NH_4_^+^ oxidation in our study, which is in line with the lower NO_2_^−^ oxidation rates (2.0-21 µmol g^−1^ day^−1^). Similar findings were obtained by Caffrey et al. (2019), whereas in other coastal sediments the rate for NO_2_^−^ oxidation exceeded that of NH_4_^+^ oxidation (Marchant et al., 2016). NH_4_^+^ oxidizers and NOB seem to occupy different niches, and are even reported to have different life strategies (Hou et al., 2018; Kitzinger et al., 2020), which could explain the observed discrepancy in NH_4_^+^ and NO_2_^−^ oxidation in different marine sediments. Although oxic conditions were present and confirmed throughout the incubations, part of the NO_2_^−^ removal in the oxic incubations was due to denitrification, as indicated by the formation of labeled N_2_O and N_2_ in the incubations to which Na^15^NO_2_ was added (treatment 6). Denitrification can occur at oxygen concentrations up to the µM range (Bonin & Raymond, 1990; Broman et al., 2021; Silvennoinen et al., 2008), and in anoxic niches associated with particles (Wan et al., 2023). Hence, the occurrence of denitrification in the oxic incubations cannot be ruled out.

Interestingly, the potential for NH_4_^+^ and NO_2_^−^ oxidation was observed for sediment sections that contain no (measurable) concentrations of oxygen *in situ*. Moreover, the relative abundance of nitrifiers and their nitrification genes was slightly higher in the anoxic deeper sediments compared to the surface sediments. NH_4_^+^ oxidizers have also been detected in other anoxic, marine environments (Du et al., 2022; Peng et al., 2013). This raises the question whether these nitrifiers are active *in situ* and if so, what role they play in these anoxic sediments. The fact that they can be revived upon the addition of NH_4_^+^ and oxygen suggests that they are metabolically active. Lipsewers et al. (2014) detected transcriptional activity from NH_4_^+^ oxidizers up to 12 cm in North Sea sediments and proposed that bioturbation may provide sufficient oxygen to sustain nitrification (Lipsewers et al., 2014). As no macrofauna were detected at Scharendijke in both winter and summer (Klomp et al., 2025), bioturbation is not expected to provide oxygen to deeper sediments at the Scharendijke basin. NH_4_^+^ oxidation activity has been observed in other anoxic marine sediments as well upon the addition of NH_4_^+^ and oxygen (Caffrey et al., 2019). More strikingly, nitrification potential in marine sediments was observed in hypoxic and even anoxic conditions (Beman et al., 2012; Bristow et al., 2016; Caffrey et al., 2019; Luo et al., 2014). Recently, oxygen production by the Thaumarchaeon *Nitrosopumilus* was reported (Hernández-Magaña & Kraft, 2024; Kraft et al., 2022), yet it is uncertain whether this also happens *in situ* and whether this could sustain nitrification in coastal sediments for prolonged times. Nonetheless, there is increasing evidence that dark oxygen production could support the widespread occurrence of aerobic metabolism in anoxic environments (Ruff et al., 2024), yet the metabolism of nitrifiers in these anoxic environments remains speculative and a topic of future studies.

### Anammox contribution to nitrogen removal is not significant in Lake Grevelingen

Anammox potential was not detected in the sediments of Lake Grevelingen. The incubations with externally added *Scalindua* biomass showed ^29^N_2_ production and thus anammox activity in both the presence and absence of sediment, indicating that the experimental set-up was suitable for detecting anammox and that the slurries were not inhibitory to anammox. Hence, the lack of ^29^N_2_ production in the incubations with sediment does indicate a lack of anammox activity in these sediments. Although these activity tests were done with sediment from 25-30 cm depth from March, anammox bacteria as well as their *hzs* genes were not detected by 16S rRNA and metagenome sequencing in any of the studied sediment sections. In contrast, anammox bacteria were detected previously in sediments up to 5 cm depth at Den Osse, another basin in Lake Grevelingen (Lipsewers et al., 2016). However, the H_2_S concentrations in the sediment at Scharendijke basin exceeded those in the sediments at Den Osse, likely inhibiting anammox (Jin et al., 2013). Moreover, the sedimentation rate at Scharendijke basin is 13 cm year^−1^ (Egger et al., 2016), making it a more dynamic system compared to Den Osse, which has a sedimentation rate up to 2 cm year^−1^ (Sulu-Gambari et al., 2018). As anammox are slow-growing bacteria, they are likely outcompeted by denitrifying microorganisms for the limited availability of NO_2_^−^ in the surface sediments of Scharendijke (Strous et al., 1998). Indeed, the formation of ^30^N_2_ upon the addition of Na^15^NO_2_ to the sediment incubations shows that NO_2_^−^ is rapidly reduced by denitrification.

### Potential for DNRA increases with NH_4_^+^ concentration at the cost of denitrification

As no anammox was detected in the sediments of Scharendijke basin, denitrification and DNRA are the main processes contributing to NO_x_ reduction in these sediments. Indeed, the genes encoding for DNRA (*nrfA/otr*) and the complete denitrification pathway (*narG/napA, nirK/nirS, norB, nosZ*) were detected at all sediment sections. Although denitrification is a modular process, the lower relative abundance of the denitrification genes compared to *gyrA* suggests that not all microorganisms possess (a) denitrification gene(s). In line with other coastal sediments, *nirS* was more abundant than *nirK* in the sediments of Lake Grevelingen (Lee & Francis, 2017; Teng et al., 2022).

The NO_3_^−^ reduction rates found in this study were lower than those reported for permeable sediments of the Wadden Sea (Gao et al., 2010) but higher than the rates reported for the sediments of the eutrophic Yangtze Estuary (Deng et al., 2015; Gao et al., 2022; Hou et al., 2013). Notably, the highest NO_3_^−^ reduction rates observed in this study were measured after 48 hrs for most sediment sections. It is therefore likely that these long incubation times could have led to the growth and enrichment of NO_3_^−^-reducing microorganisms (Massana et al., 2001), and that the rates presented here are thus an overestimation of the *in situ* NO_3_^−^ reduction rates. Indeed the incubations led to the enrichment of certain microbial groups compared to the original sediment. Moreover, NO_3_^−^ reduction rates can vary greatly, depending on a complex interplay between geochemical conditions such as salinity, organic carbon and sulfide concentrations and the microbial community composition (Deng et al., 2015; Gao et al., 2022; Li et al., 2025).

In both seasons and at all depths studied, denitrification dominated NO_3_^−^ reduction, exceeding the contribution of DNRA. In contrast to the NH_4_^+^ oxidation potential, NO_3_^−^ reduction displayed redox zone-related patterns. Consistent with previous findings, the DNRA contribution increased with depth, where sediments are more reducing and exhibit an electron donor surplus (Fan et al., 2024; Jäntti & Hietanen, 2012). As DNRA can utilize more electrons than denitrification, these reduced zones likely favor DNRA (Tiedje et al., 1983). A higher contribution of DNRA was also observed in incubations amended with electron donors such as MMA (treatment 11). Environmental factors, such as the C/N ratio and NO_3_^−^ versus NO_2_^−^ supplementation, influence the competition between denitrification and DNRA (Behrendt et al., 2014; Brin et al., 2015; Jia et al., 2020; Kraft et al., 2014; Van Den Berg et al., 2017; Yoon et al., 2015). This likely also affects the competition between denitrification and DNRA in the treatment 9 bottles. Sulfide can stimulate DNRA (Murphy et al., 2020), contributing to the higher relative contribution of DNRA in the deeper, sulfidic sediments from March. While sulfide can act as an electron donor for both denitrification and DNRA, it may also inhibit the last step of denitrification (Brunet & Garcia-Gil, 1996). Similarly, sulfide inhibited the anaerobic oxidation of methane in the eutrophic sediments of the Stockholm Archipelago (Dalcin Martins et al., 2024). As sulfide is present at mM concentrations in the deeper sediments in March, the effect of sulfide on the denitrification to DNRA ratio in sediments from Lake Grevelingen cannot be ruled out (Klomp et al., 2025; Van Helmond et al., 2025; Żygadłowska et al., 2023).

Unexpectedly, however, the contribution of DNRA to NO_3_^−^ reduction was lower in surface sediments of September than the deeper sediments of March, despite the fact that both were anoxic and sulfidic. Concomitant with this finding, the NO_3_^−^-amended incubations with sediment from March and September resulted in the enrichment of different bacterial communities. The NO_3_^−^-amended incubations with sediment from March were dominated by the bacterial families *Arcobacteraceae* in the surface sediments and *Sulfurimonadaceae* and *Desulfuromonadaceae* in the deeper, more reduced and sulfidic sediments. *Arcobacteraceae* has been proposed to play a role in both denitrification and DNRA (Isokpehi et al., 2024; Kraft et al., 2014; Li et al., 2024). Some *Sulfurimonadaceae* genera can reduce NO_3_^−^ coupled to sulfur oxidation and possess the genes for denitrification (Bruckner et al., 2013; Cai et al., 2014; Grote et al., 2012; Takai et al., 2006), and the sulfate-reducing family *Desulfuromonadaceae* has been detected in incubations with high sulfide concentrations in which DNRA dominated over denitrification (Murphy et al., 2020). Potentially, *Desulfurmonadaceae* could reduce the oxidized sulfur produced upon sulfide oxidation by *Sulfurimonadaceae*. In contrast, the NO_3_^−^-amended incubations with surface sediment from September were dominated by *Nitrincolaceae*. This bacterial family is often detected following phytoplankton blooms, but their role in NO_3_^−^ reduction remains elusive (Thiele et al., 2023). Interestingly, *Ca*. Methylomirabilis and *Methanoperedenaceae* were not detected in any sediment samples, suggesting that nitrate-dependent anaerobic methane oxidation (Ettwig et al., 2010; Haroon et al., 2013) does not play a significant role in the sediments of Scharendijke basin, despite the high CH_4_ concentrations in these sediments (Żygadłowska et al., 2023). The balance between denitrification and DNRA not only depends on the geochemistry of the sediment but also on the tight interplay between the chemical parameters and the microbial community (Jäntti et al., 2022; Kraft et al., 2014; Neubauer et al., 2019).

Interestingly, MMA was consumed in all incubations to which it was added and stimulated denitrification. MMA is a well-studied (marine) substrate for methylotrophic methanogenesis (Krause & Treude, 2021; Lynes et al., 2024), but only a few studies have reported on bacteria capable of oxidizing MMA anaerobically with NO_3_^−^ (Kim et al., 2001). MMA has been detected in marine sediments in nM to µM range (Fitzsimons et al., 2023) and could thus play a significant role in NO_3_^−^ reduction *in situ*, either directly or indirectly. The effect of MMA addition on denitrification was most apparent in the NO_3_^−^ and MMA-amended bottles with surface sediment from March, where complete reduction of NO_3_^−^ and NO_2_^−^ was observed in the presence but not in the absence of MMA.

The maximum potential reduction rate of NO_x_ generally exceeded that of N_2_O reduction, which indeed led to the accumulation of some N_2_O in the NO_x_-amended incubations (treatments 8, 9). *In situ* denitrification of NO_3_^−^ could thus lead to N_2_O emissions from the sediments of Lake Grevelingen, as has been observed for Wadden Sea sediments (Marchant et al., 2018). In this system, Flavobacteriia were the dominant N_2_O reducers, proposedly acting as N_2_O-reducing specialists possessing *nosZ* as sole denitrification gene (Marchant et al., 2018). Notably, the majority of the detected *nosZ* sequences in the sediments from Lake Grevelingen were classified as ‘Sec-nosZ’, and they were affiliated with Flavobacteriia as well. NosZ proteins with a Sec signal peptide belong to the atypical clade II NosZ, which is commonly found in microorganisms lacking other denitrification genes (Jones et al., 2013; Sanford et al., 2012).

The question remains to what extent denitrification and DNRA play a role in Lake Grevelingen sediments *in situ*, despite the observed high potential for these processes. Both require NO_3_^−^ or NO_2_^−^, which were below the detection limit in the sediment. Anoxia, such as in the deeper sediments of March and the surface sediments of September, is expected to inhibit NO_x_ supply from nitrification and hence lead to lower denitrification rates (Childs et al., 2002; Jäntti & Hietanen, 2012). Moreover, the accumulation of sulfide in these sediments can favor processes such as DNRA while it can inhibit other processes, such as anammox and denitrification. Nonetheless, this study shows that there is good potential for nitrification, denitrification, and DNRA in Lake Grevelingen sediments and that these processes can be rapidly reactivated upon the addition of the required substrates. These processes likely contribute to the removal of nitrogen and further depletion of electron acceptors in the surface sediments, yet their *in situ* relevance at depth in the sediment at Scharendijke requires further investigation.

## Conclusion

We investigated the microbial nitrogen cycling potential in sediments from seasonally euxinic, coastal Lake Grevelingen by combining incubation experiments with 16S rRNA and metagenome sequencing and linked the results to the geochemical context of the ecosystem. We showed that the potential for NH_4_^+^ oxidation and the presence of a nitrifying microbial community were not restricted to the oxic surface sediments in winter, as these were also detected in anoxic sediments. Likewise, the potential for anaerobic reduction of NO_3_^−^, NO_2_^−^ and the potent greenhouse gas N_2_O was observed across sediment depths and seasons. Anammox was not detected, likely because of the slow growth of anammox bacteria and their sensitivity to sulfide, and NO_x_ reduction therefore proceeded mainly through denitrification, followed by DNRA. As the sediments contain low to no measurable amounts of the required electron acceptors, the *in situ* relevance of the studied processes requires future investigation. Combined, our findings highlight the potential for removal of nitrogen from eutrophic coastal systems, providing valuable insights that are of relevance for the management of coastal environments under increasing anthropogenic pressures.

## Acknowledgements

This research was supported by ERC Synergy Grant MARIX 854088. We thank all the MARIX team members, the crew of R/V Navicula of Royal NIOZ and all participants on the cruises.

