## Supplementary data for "Sediments from a seasonally euxinic coastal ecosystem show high nitrogen cycling potential"

Appendix

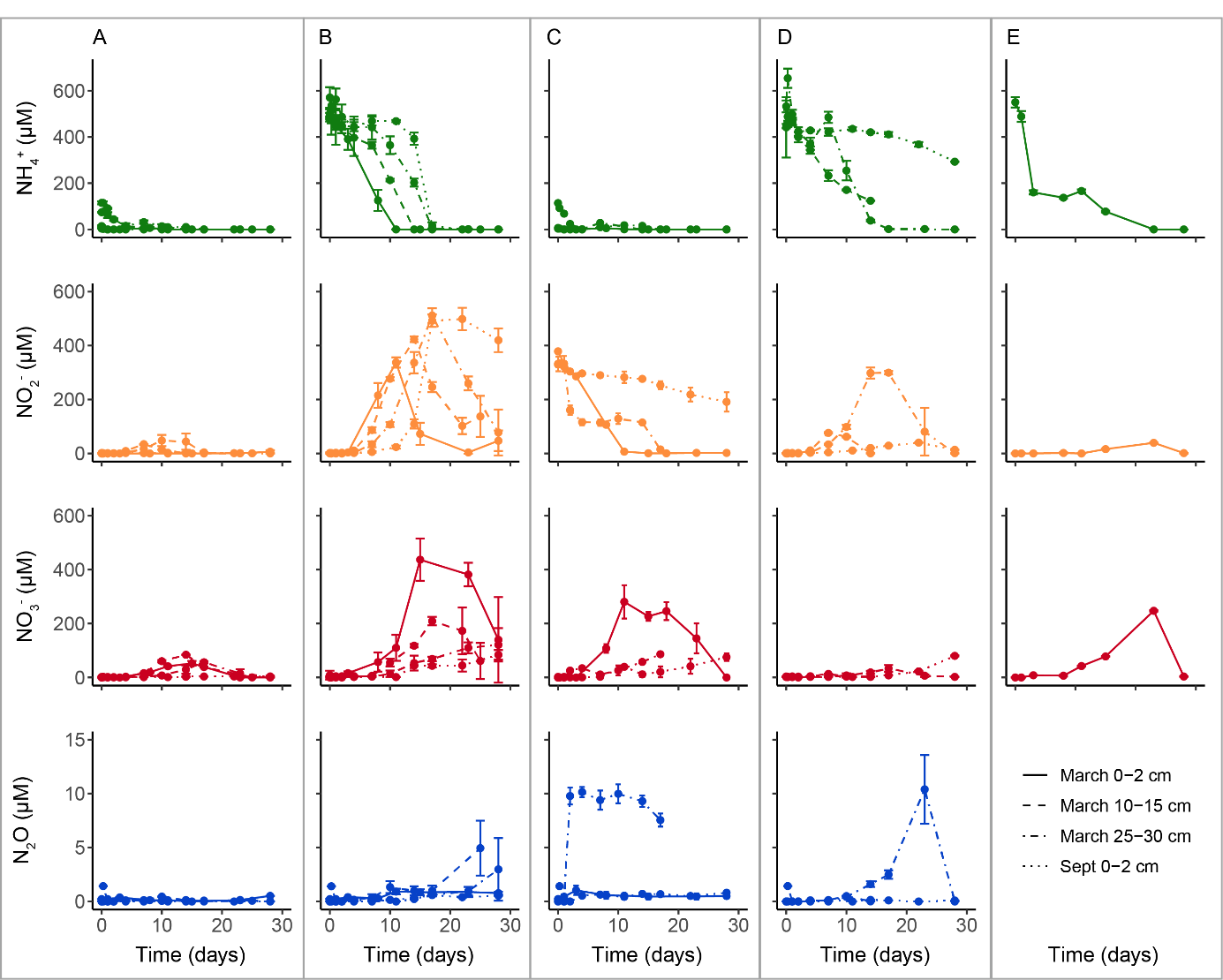

Figure S1. Concentrations of dissolved NH_4_^+^ (green), NO_2_^-^ (orange), NO_3_^-^ (red), and headspace N_2_O (blue) over time in oxic incubations with sediment from Lake Grevelingen from March 2023 10-15 and 25-30 cm depth, September 2023 0-2 cm depth, and March 2025 0-2 cm depth, supplemented with (A) no substrate (treatment 1), (B) 0.5 mM NH_4_Cl (treatment 2), (C) 0.25 mM NaNO2 (treatment 3), (D) 0.5 mM NH_4_Cl +2% CH_4_ (treatment 4) and (E) 0.5 mM NH_4_Cl in Erlenmeyers (treatment 5). Error bars indicate the standard deviation of the triplicate measurements for each sediment-treatment combination.

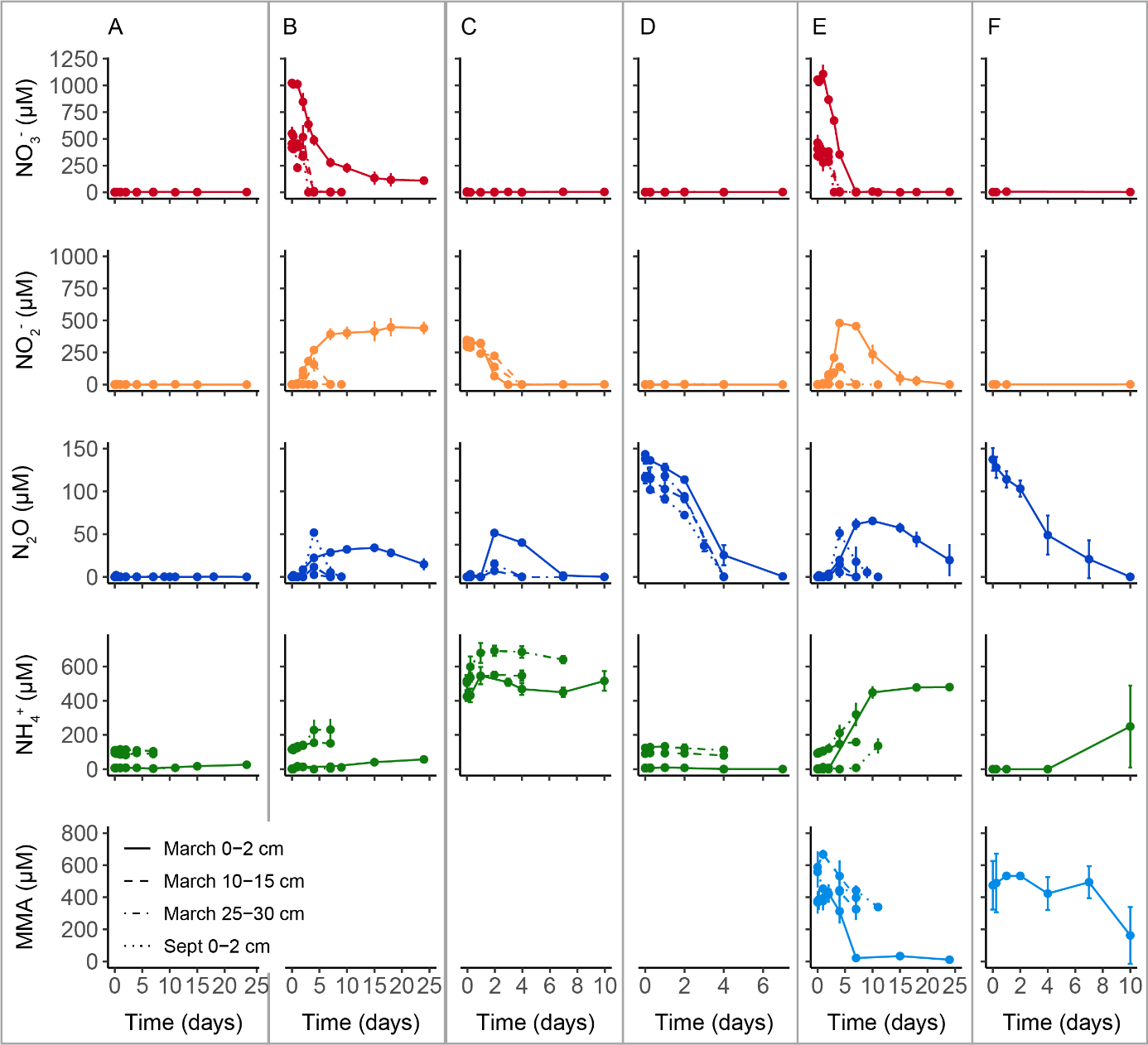

Figure S2. Concentrations of dissolved NO_3_^-^ (red), NO_2_^-^ (orange), NH_4_^+^ (green), MMA (light blue) and headspace N_2_O (blue) over time in anoxic incubations with sediment from Lake Grevelingen from March 2023, 0-2, 10-15 and 25-30 cm depth, and September 2023, 0-2 cm depth. Bottles were supplemented with (A) no substrate (treatment 7), (B) 0.5-1 mM NaNO_3_ (treatment 8), (C) 0.25 mM NaNO_2_ + 0.5 mM NH_4_Cl + 2% CO_2_ (treatment 9), (D) 0.5 % N_2_O (treatment 10), (E) 0.5-1 mM NaNO_3_ + 0.5 mM MMA (treatment 11), and (F) 0.5% N_2_O + 0.5 mM MMA (treatment 12). Error bars indicate the standard deviation of the triplicate measurements for each sediment-treatment combination.

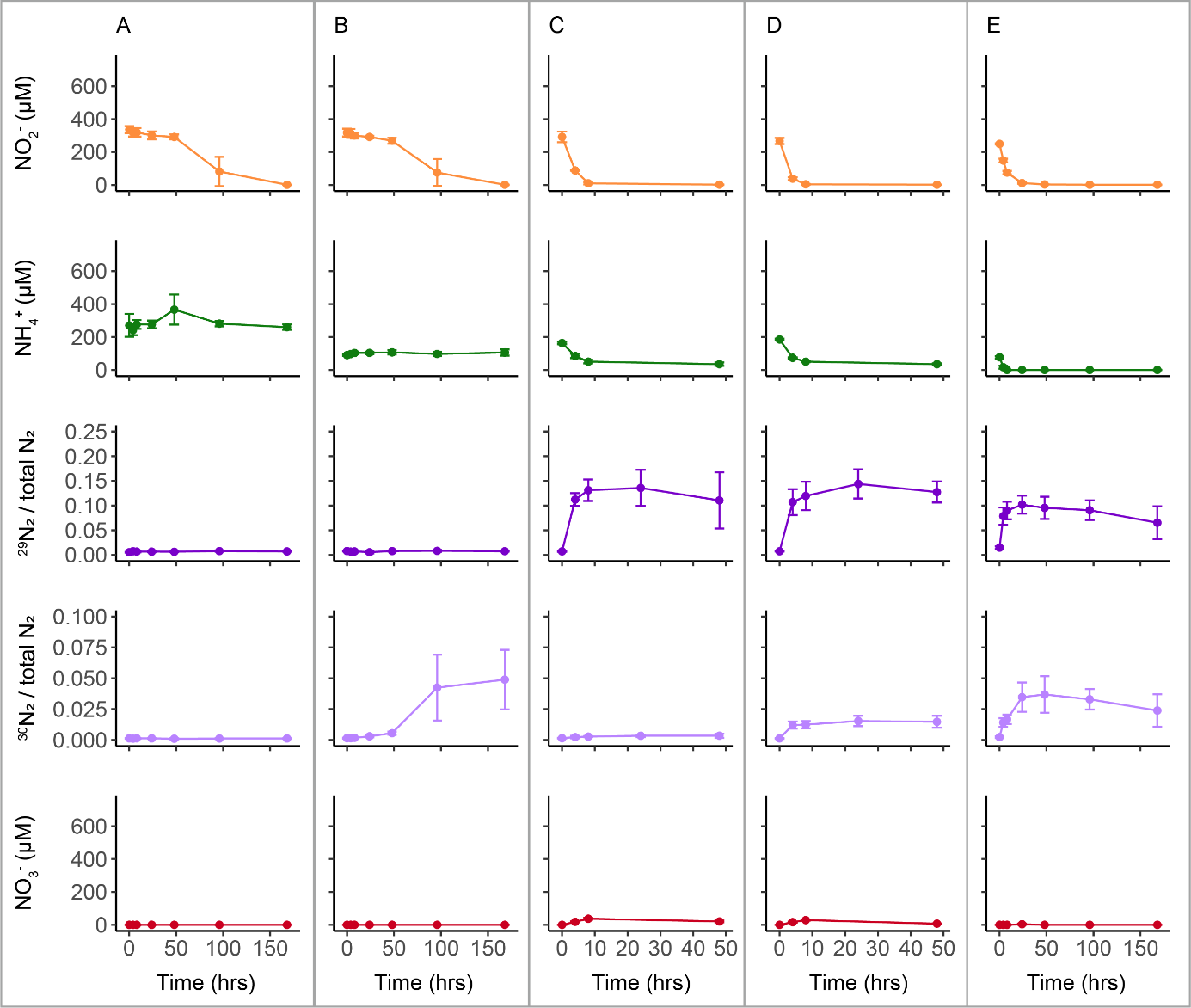

Figure S3. Concentrations of dissolved NO_2_^-^ (orange), NH_4_^+^ (green) and NO_3_^-^ (red) and the ratios of ^29^N_2_ and ^30^N_2_ over total N_2_ (dark blue and light blue, respectively) over time in incubations with various inoculums and amendments:(A) sediment from 25-30 cm from March 2024 amended with 0.2 mM NaNO_2_ and ^15^NH_4_Cl (treatment 13), (B) sediment from 25-30 cm from March 2024 amended with 0.2 mM Na^15^NO_2_ (treatment 14), (C) Scalindua biomass amended with 0.2 mM NaNO_2_ and ^15^NH_4_Cl (treatment 15), (D) Scalindua biomass amended with 0.2 mM Na^15^NO_2_ and NH_4_Cl (treatment 16), (E) sediment from 25-30 cm from March 2024 and Scalindua biomass amended with 0.2 mM Na^15^NO_2_ (treatment 17). Error bars indicate the standard deviation of the triplicate measurements for each sediment-treatment combination.

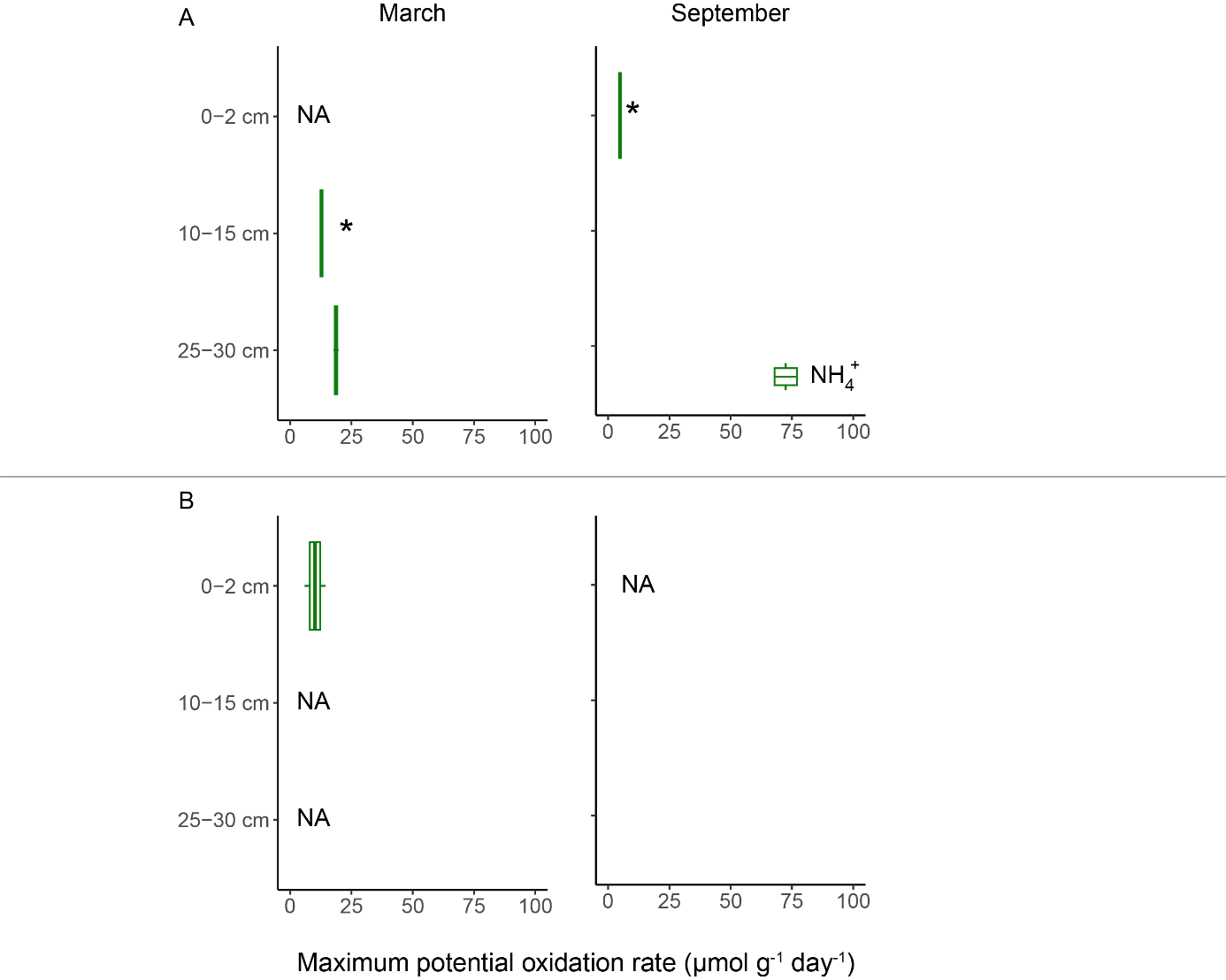

Figure S 4. Maximum potential rates (µmol g^-1^ day^-1^) of NH_4_^+^ oxidation at different sediment depths in March and September (left and right panel, respectively), as measured in duplicate incubations amended with 0.5 mM NH_4_Cl in the presence of (A) 2% CH_4_ (treatment 4) and (B) atmospheric oxygen (treatment 5). NA: treatment not included for incubations with this sediment section; *: single bottle included for this treatment instead of duplicate bottles.

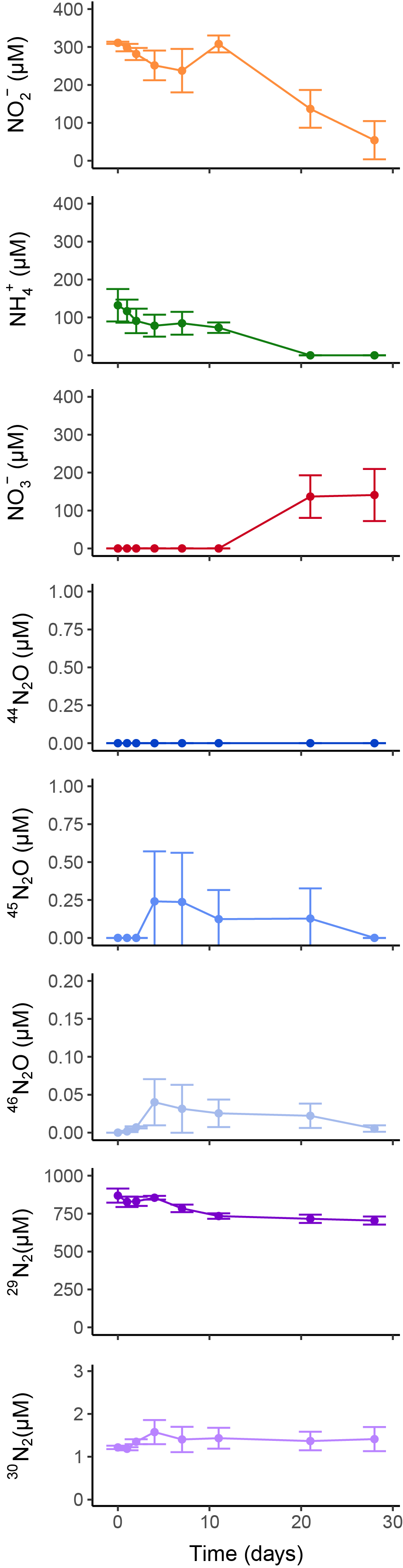

Figure S5. Concentrations of dissolved NO_3_^-^ (red), NO_2_^-^ (orange), NH_4_^+^ (green), and the headspace concentrations of N_2_O (blue) and N_2_ (purple) isotopes over time in oxic bottle incubations with sediment from Lake Grevelingen from March 2024 25-30 cm depth. Bottles were supplemented with 0.2 mM Na^15^NO_2_ (treatment 6). Error bars indicate the standard deviation of the triplicate measurements for each sediment-treatment combination.

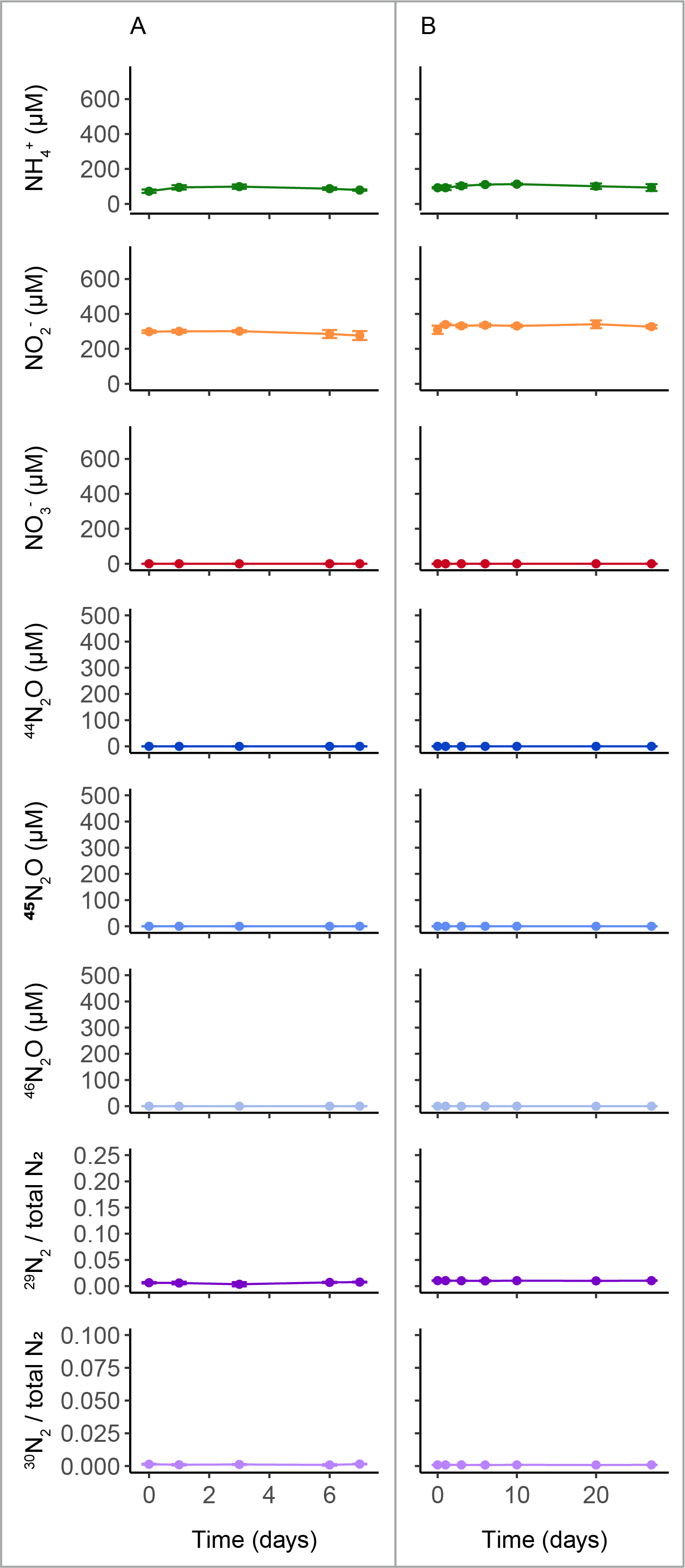

Figure S6. Concentrations of dissolved NH_4_^+^ (green), NO_2_^-^ (orange), NO_3_^-^ (red), and the headspace concentrations of N_2_O (blue) and N_2_ (purple) isotopes over time in autoclaved oxic (A; treatment 18) and anoxic (B; treatment 19) incubation bottles inoculated with sediment from 25-30 cm depth from March 2024, supplemented with 0.2 mM Na^15^NO_2_. Error bars indicate the standard deviation of the triplicate measurements for each sediment-treatment combination.

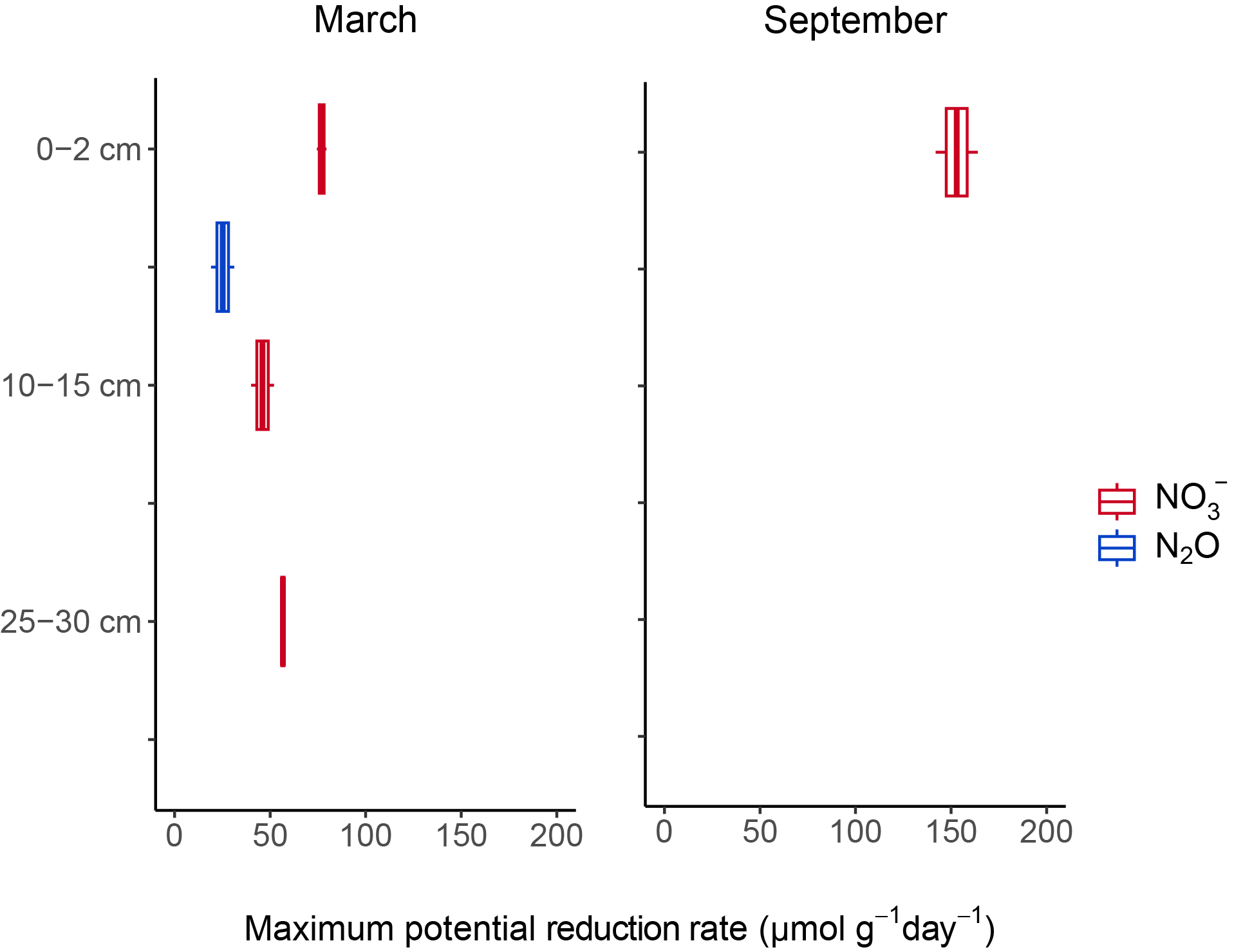

Figure S7. Maximum potential rates (µmol g^-1^ day^-1^) of NO_3_^-^ (red) and N_2_O (blue) oxidation at different sediment depths in March and September (left and right panel, respectively), as measured in duplicate incubation bottles amended with 0.5-1 mM NaNO_3_ or 0.5% N_2_O in presence of 0.5 mM MMA (treatments 11 and 12, respectively). NA: treatment not included for incubations with this sediment section.

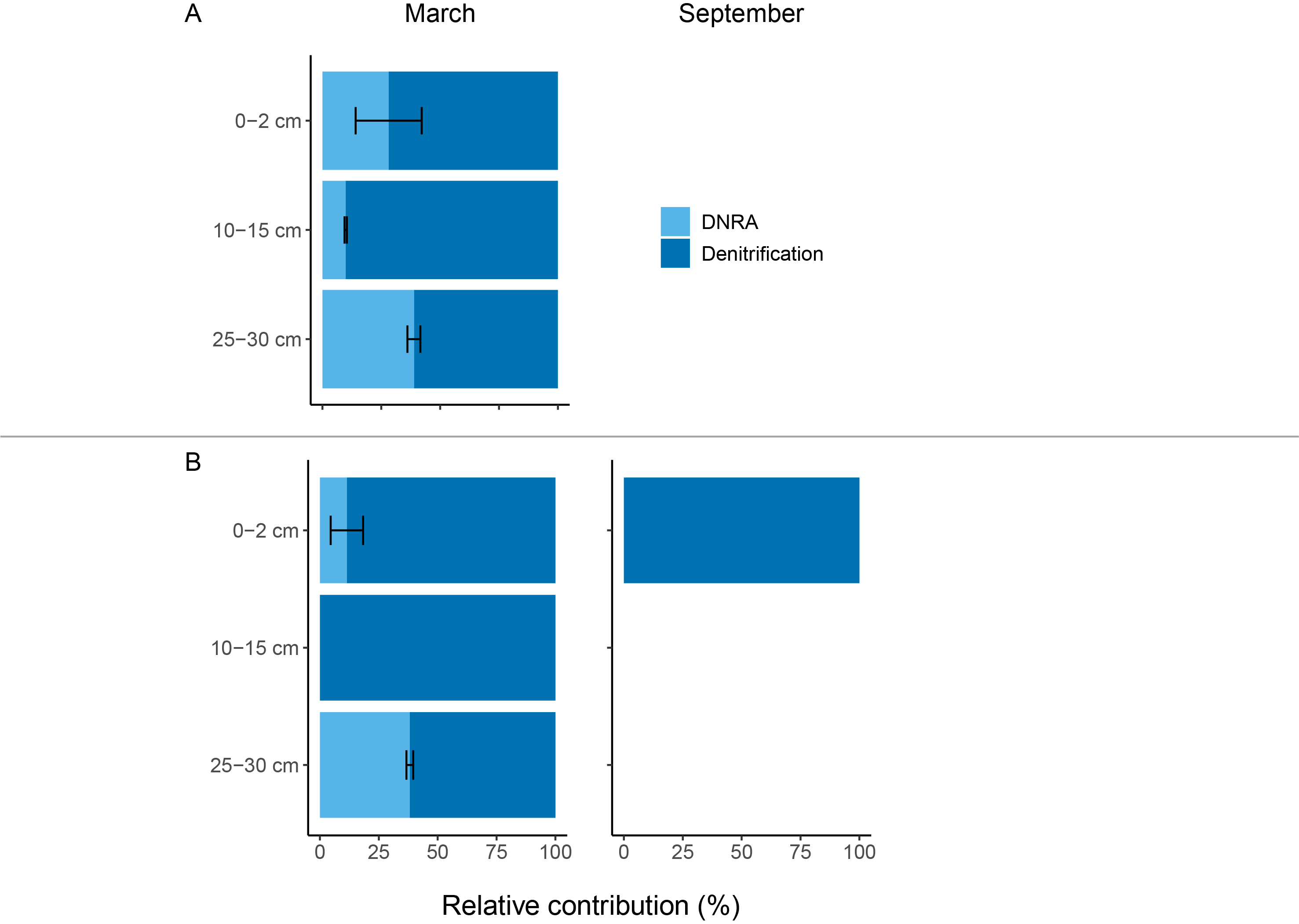

*Figure S8. Relative contribution of DNRA and denitrification at different sediment depths in March and September left and right panel, respectively) to (A) NO_2_^-^ reduction in presence of excess NH_4_^+^, as measured in incubation bottles supplemented with 0.25 mM NaNO_2_ and 0.5 mM NH_4_Cl (treatment 9), and (B) NO_3_^-^ reduction in presence of MMA, as measured in incubation bottles supplemented with 0.5-1 mM NaNO_3_ and 0.5 mM MMA (treatment 11). Error bars indicate the standard deviation between duplicate bottles.*

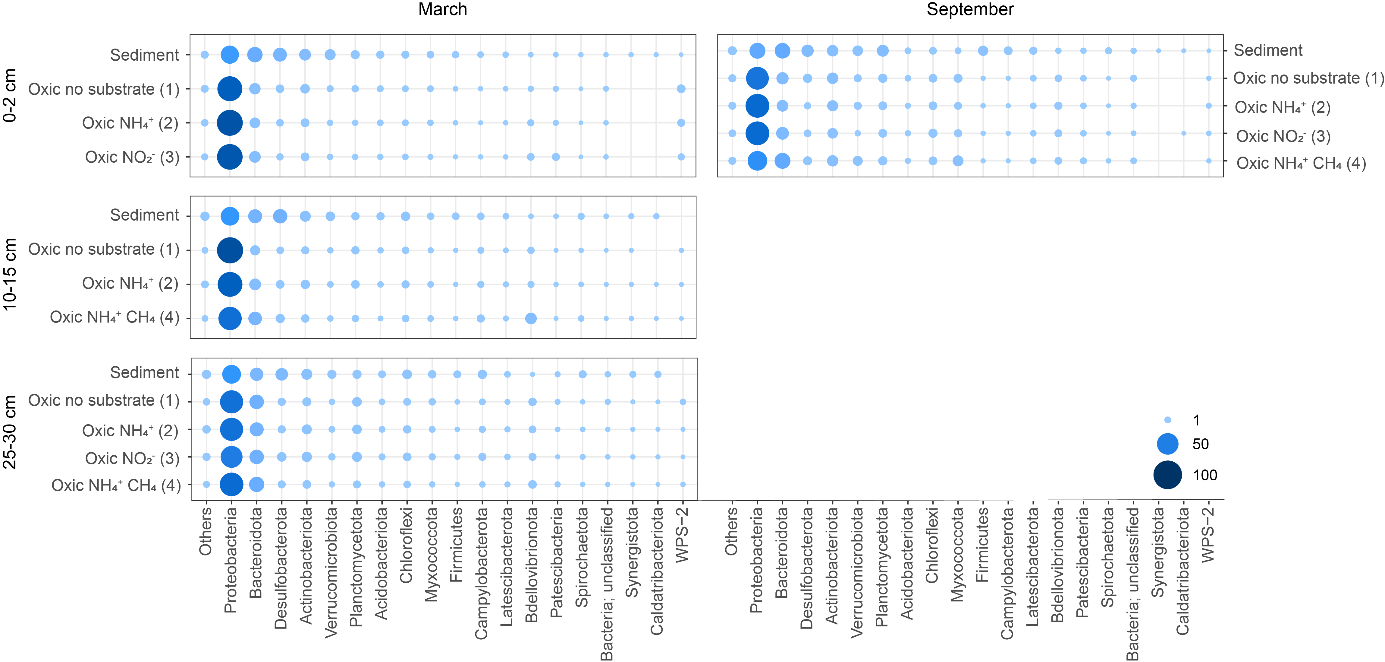

Figure S9. Relative abundances of the bacterial phyla in the original sediment and the oxic incubations set up with each sediment section (left panel: March, right panel: September). Phyla present < 1% in all presented samples have been grouped in ‘Others’. The numbers in between brackets refer to the number of the corresponding treatment.

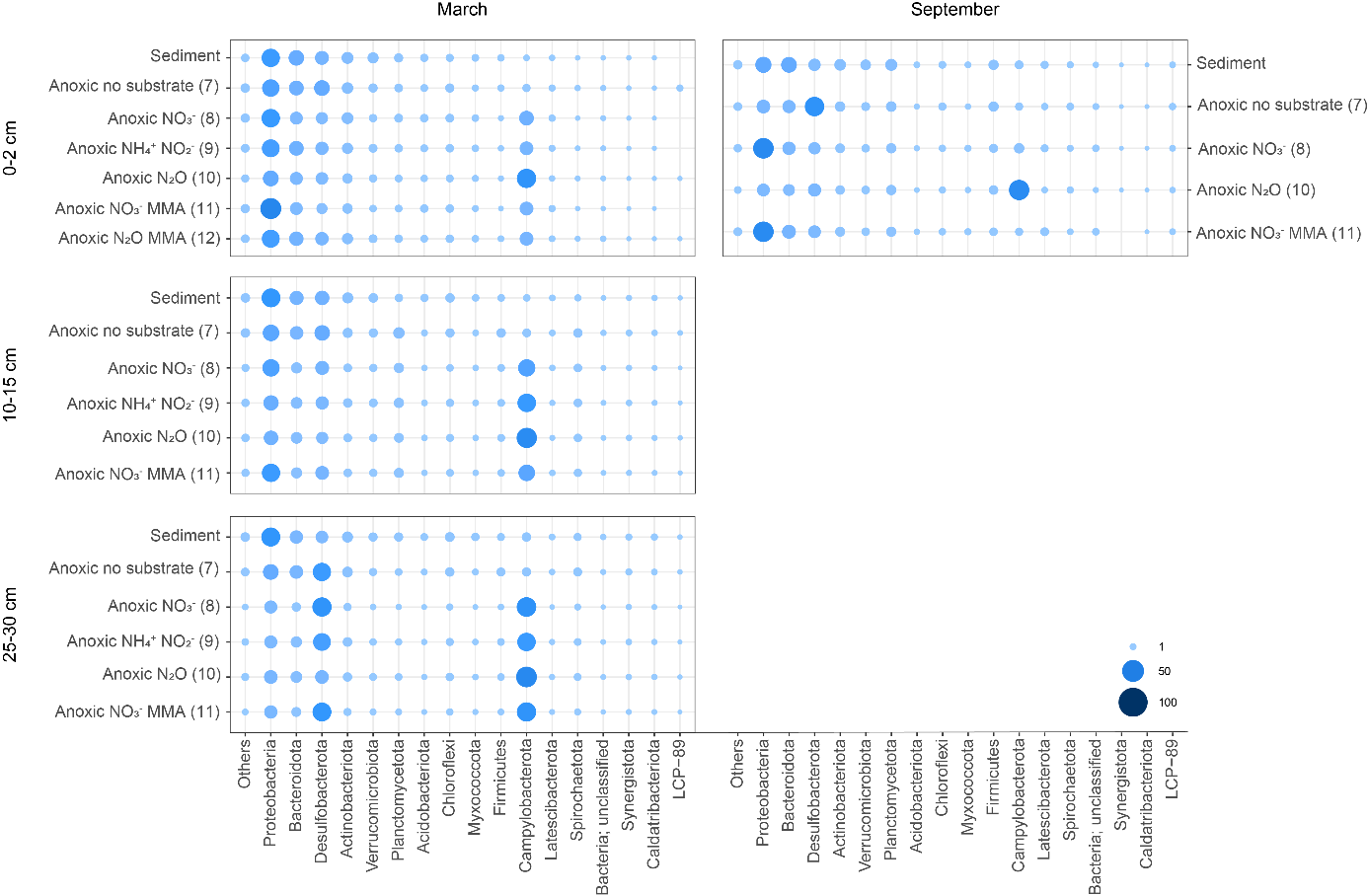

Figure S10. Relative abundances of the bacterial phyla in the original sediment and the anoxic incubations set up with each sediment section (left panel: March, right panel: September). Phyla detected < 1% in all presented samples have been grouped in ‘Others’. The numbers in between brackets refer to the number of the corresponding treatment.

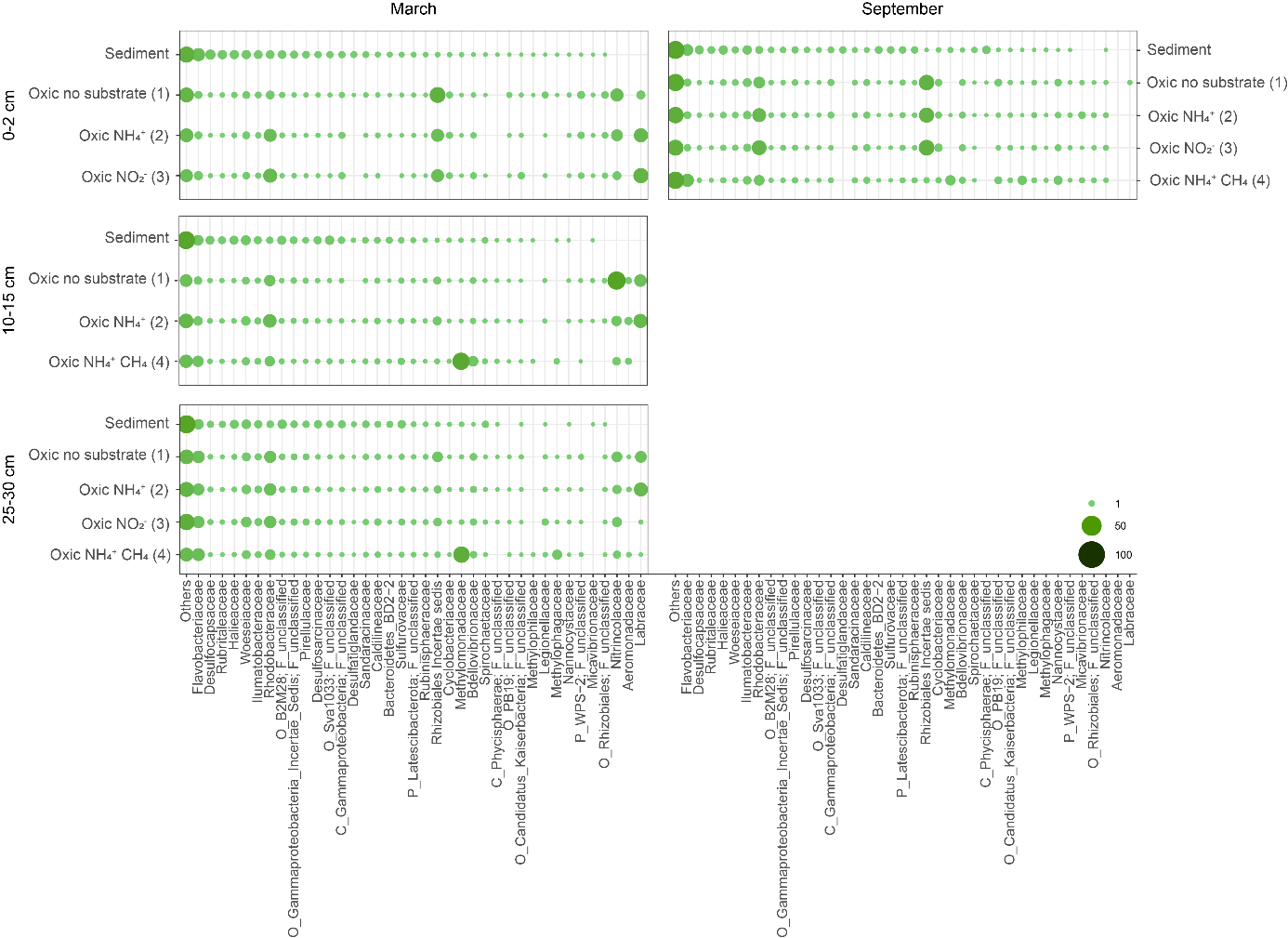
Figure S11. Relative abundances of the bacterial families in the original sediment and the oxic incubations set up with each sediment section (left panel: March, right panel: September). Families detected < 2% in all presented samples have been grouped in ‘Others’. The numbers in between brackets refer to the number of the corresponding treatment.

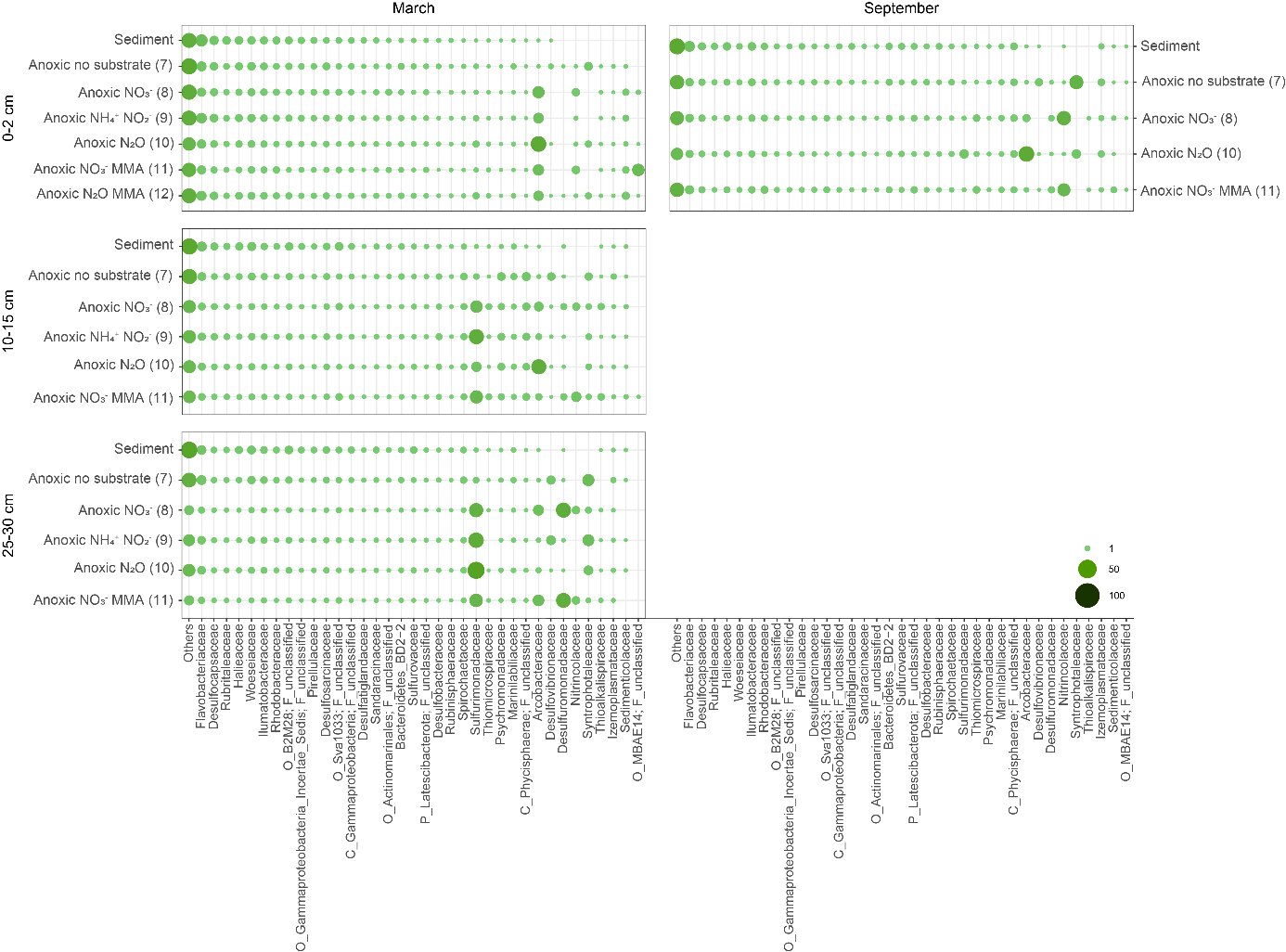

Figure S12. Relative abundances of the bacterial families in the original sediment and the anoxic incubations set up with each sediment section (left panel: March, right panel: September). Families detected < 2% in all presented samples have been grouped in ‘Others’. The numbers in between brackets refer to the number of the corresponding treatment.

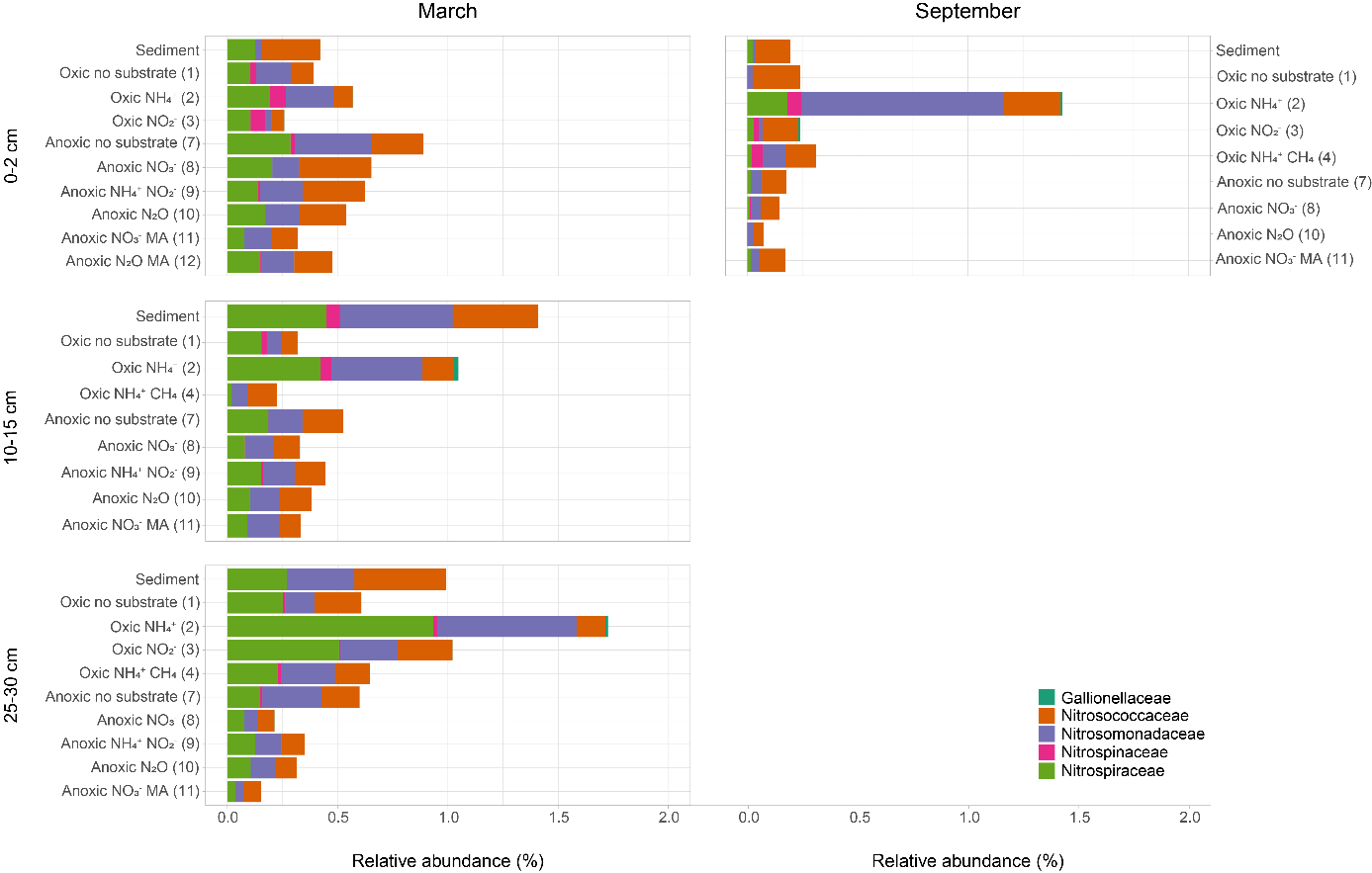

Figure S13. Relative abundances of nitrifying bacteria at the family level, grouped per sediment section (left panel: March, right panel: September). The numbers in between brackets refer to the number of the corresponding treatment.

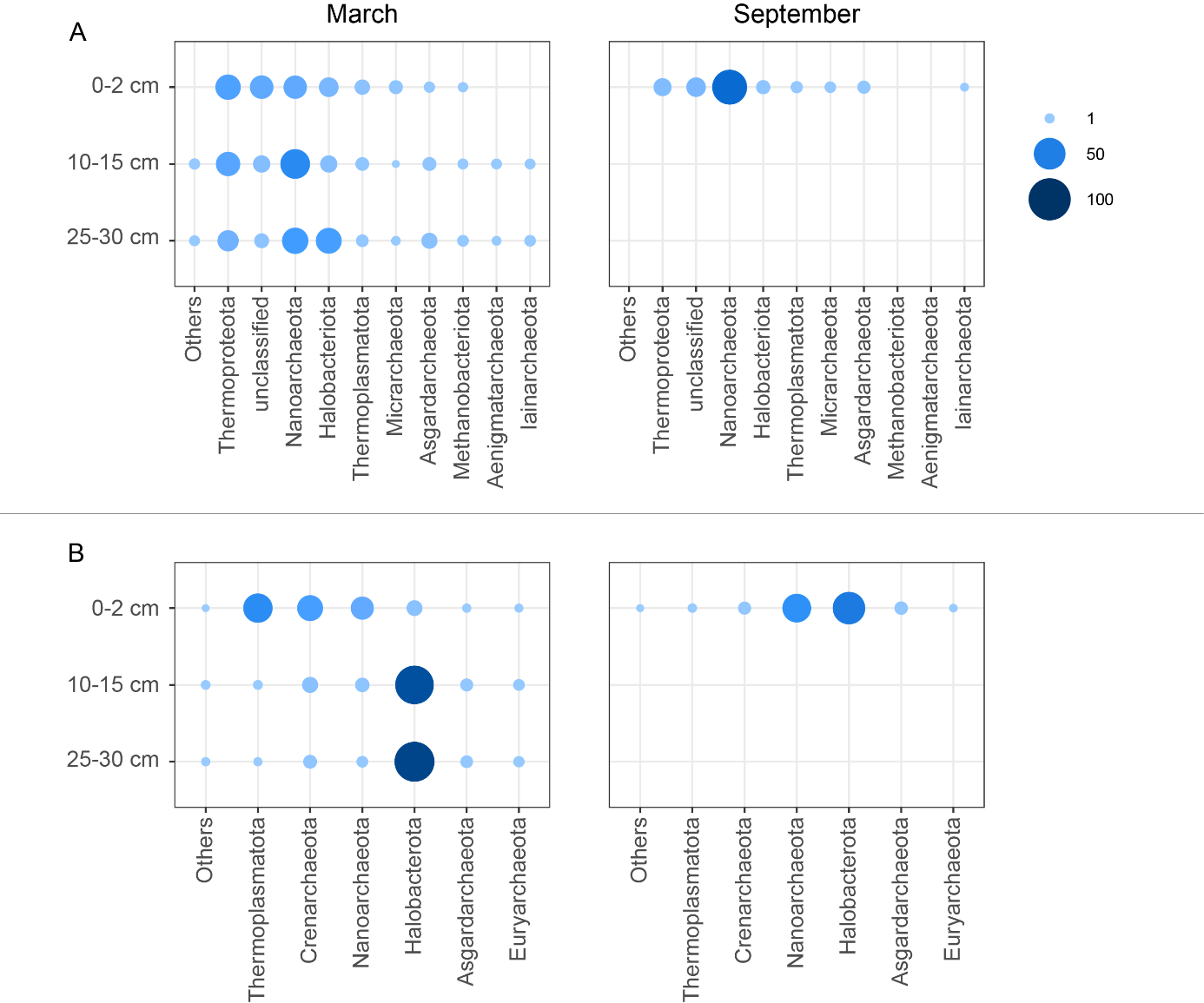

Figure S14. Relative abundances of archaeal phyla in March (left panel) and September (right panel). SingleM analysis of the metagenome data revealed the abundances in the original sediment at the four studied depths and seasons (A), whereas 16S rRNA amplicon sequencing was done to identify the relative abundances in the oxic incubations amended with 0.5 mM NH_4_Cl (treatment 2) (B). Phyla present < 1% in all presented samples have been grouped in ‘Others’.

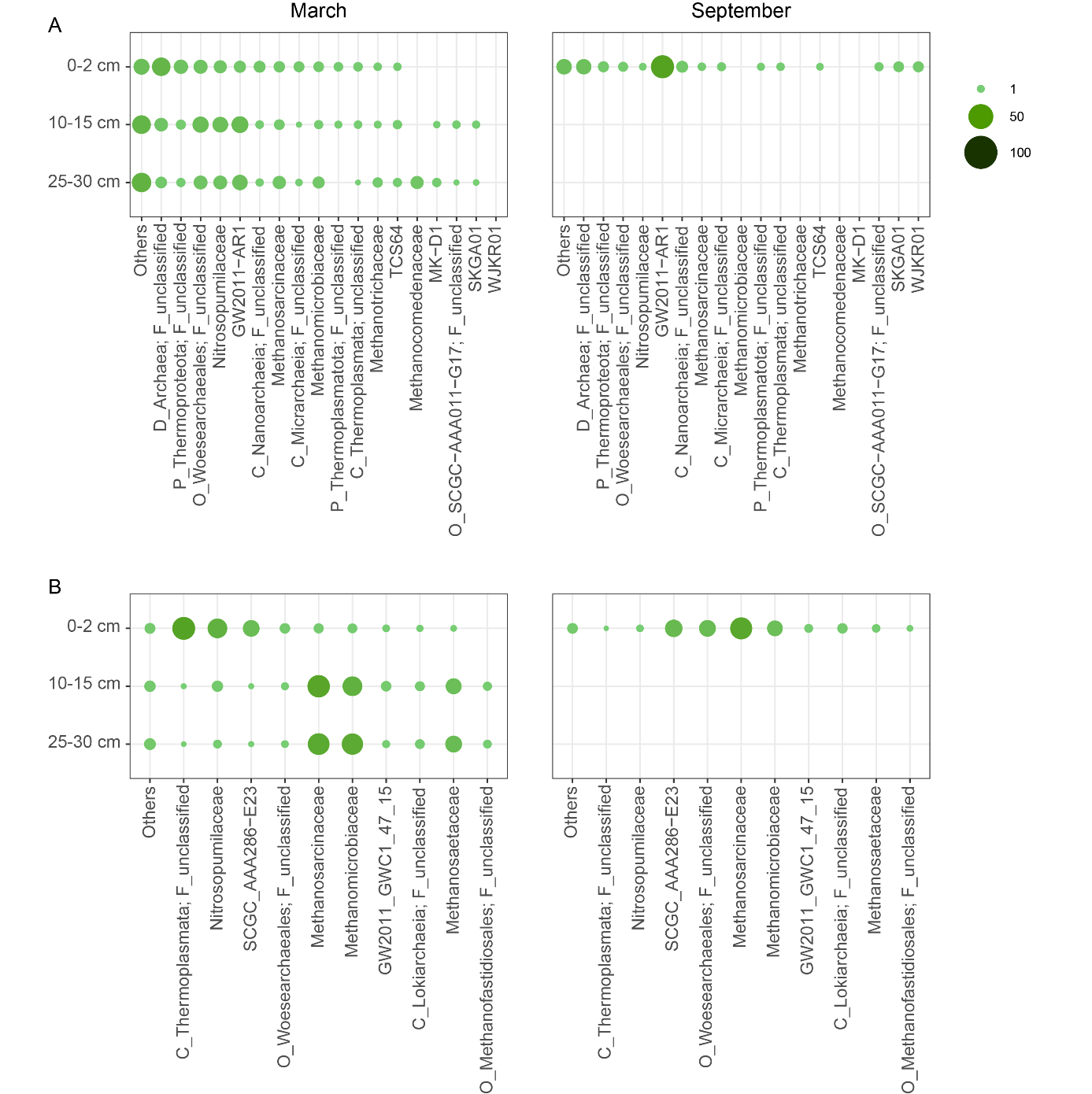

Figure S15. Relative abundances of archaeal families in March (left panel) and September (right panel). SingleM analysis of the metagenome data revealed the abundances in the original sediment at the four studied depths and seasons (A), whereas 16S rRNA amplicon sequencing was done to identify the relative abundances in the oxic incubations amended with 0.5 mM NH_4_Cl (treatment 2) (B). Families present < 2% in all presented samples have been grouped in ‘Others’.

Table S1. The bottle incubations set up for studying the nitrogen cycling potential in sediments from Lake Grevelingen.

| Treatment number | Inoculum | Treatment |
| --- | --- | --- |
| 1 | Sediment March 2023 10-15 cm  Sediment March 2023 25-30 cm  Sediment Sept 2023 0-2 cm  Sediment March 2024 0-2 cm | Oxic, no substrates |
| 2 | Sediment March 2023 10-15 cm  Sediment March 2023 25-30 cm  Sediment Sept 2023 0-2 cm  Sediment March 2024 0-2 cm | Oxic, 0.5 mM NH_4_Cl |
| 3 | Sediment March 2023 25-30 cm  Sediment Sept 2023 0-2 cm  Sediment March 2024 0-2 cm | Oxic, 0.25 mM NaNO_2_ |
| 4 | Sediment March 2023 10-15 cm  Sediment March 2023 25-30 cm  Sediment Sept 2023 0-2 cm | Oxic, 0.5 mM NH_4_Cl + 2% CH_4_ |
| 5 | Sediment March 2024 0-2 cm | Oxic, 0.5 mM NH_4_Cl (Erlenmeyers) |
| 6 | Sediment March 2024 25-30 cm | Oxic, 0.2 mM Na^15^NO_2_ |
| 7 | Sediment March 2023 0-2 cm  Sediment March 2023 10-15 cm  Sediment March 2023 25-30 cm  Sediment Sept 2023 0-2 cm | Anoxic, no substrates |
| 8 | Sediment March 2023 0-2 cm  Sediment March 2023 10-15 cm  Sediment March 2023 25-30 cm  Sediment Sept 2023 0-2 cm | Anoxic, 0.5-1 mM NaNO_3_ |
| 9 | Sediment March 2023 0-2 cm  Sediment March 2023 10-15 cm  Sediment March 2023 25-30 cm | Anoxic, 0.25 mM NaNO_2_ + 0.5 mM NH_4_Cl + 2% CO_2_ |
| 10 | Sediment March 2023 0-2 cm  Sediment March 2023 10-15 cm  Sediment March 2023 25-30 cm  Sediment Sept 2023 0-2 cm | Anoxic, 0.5% N_2_O |
| 11 | Sediment March 2023 0-2 cm  Sediment March 2023 10-15 cm  Sediment March 2023 25-30 cm  Sediment Sept 2023 0-2 cm | Anoxic, 0.5-1 mM NaNO_3_ + 0.5 mM MMA |
| 12 | Sediment March 2023 0-2 cm | Anoxic, 0.5% N_2_O + 0.5 mM MMA |
| 13 | Sediment March 2024 25-30 cm | Anoxic, 0.2 mM Na^14^NO_2_ + 0.2 mM ^15^NH_4_Cl |
| 14 | Sediment March 2024 25-30 cm | Anoxic, 0.2 mM Na^15^NO_2_ |
| 15 | *Scalindua* biomass | Anoxic, 0.2 mM Na^14^NO_2_ + 0.2 mM ^15^NH_4_Cl |
| 16 | *Scalindua* biomass | Anoxic, 0.2 mM Na^15^NO_2_ + 0.2 mM ^14^NH_4_Cl |
| 17 | Sediment March 2024 25-30 cm + *Scalindua* biomass | Anoxic, 0.2 mM Na^15^NO_2_ |
| 18 | Sediment March 2024 25-30 cm | Oxic, autoclaved, 0.2 mM Na^15^NO_2_ |
| 19 | Sediment March 2024 25-30 cm | Anoxic, autoclaved, 0.2 mM Na^15^NO_2_ |

Table S2. Mass balance of N in oxic incubation bottles with sediment from 25-30 cm depth from March 2024, amended with 0.2 mM Na^15^NO_2_ (treatment 6), calculated between the start and end of the incubation period (day 0 and day 28, respectively) for replicate bottles 1 and 2.

|  | Treatment 6, day 0 | | | | Treatment 6, day 28 | | | |
| --- | --- | --- | --- | --- | --- | --- | --- | --- |
|  | Amount of compound (µmol) | | Total amount of N per compound (µmol) | | Amount of compound left (µmol) | | Total amount of N per compound (µmol) | |
|  | Bottle 1 | Bottle 2 | Bottle 1 | Bottle 2 | Bottle 1 | Bottle 2 | Bottle 1 | Bottle 2 |
| NO_3_^-^ | 0 | 0 | 0 | 0 | 3.2 | 8.0 | 3.2 | 8.0 |
| NO_2_^-^ | 15.5 | 15.7 | 15.5 | 15.7 | 4.0 | 0.32 | 4.0 | 0.32 |
| NH_4_^+^ | 8.6 | 4.7 | 8.6 | 4.7 | 0 | 0 | 0 | 0 |
| ^44^N_2_O | 0 | 0 | 0 | 0 | 0 | 0 | 0 | 0 |
| ^45^N_2_O | 0 | 0 | 0 | 0 | 0 | 0 | 0 | 0 |
| ^46^N_2_O | 0 | 0 | 0 | 0 | 0.00075 | 0.00012 | 0.0015 | 0.00024 |
| ^28^N_2_ | 5454.8 | 5506.1 | 10909.7 | 11012.2 | 5046.3 | 4788.8 | 10092.5 | 9577.6 |
| ^29^N_2_ | 61.4 | 60.6 | 122.7 | 121.3 | 57.2 | 56.4 | 114.4 | 112.7 |
| ^30^N_2_ | 0.086 | 0.086 | 0.17 | 0.17 | 0.14 | 0.094 | 0.27 | 0.19 |
| NO_3_^-^ taken out | - | - | - | - | 0.13 | 0.28 | 0.13 | 0.28 |
| NO_2_^-^ taken out | - | - | - | - | 2.6 | 2.9 | 2.6 | 2.9 |
| NH_4_^+^ taken out | - | - | - | - | 1.1 | 0.62 | 1.1 | 0.62 |
| ^44^N_2_O taken out | - | - | - | - | 0 | 0 | 0 | 0 |
| ^45^N_2_O taken out | - | - | - | - | 0.00057 | 0 | 0.0011 | 0 |
| ^46^N_2_O taken out | - | - | - | - | 0.000077 | 0.000017 | 0.00015 | 0.000033 |
| ^28^N_2_ taken out | - | - | - | - | 157.5 | 157.5 | 315.0 | 315.0 |
| ^29^N_2_ taken out | - | - | - | - | 1.8 | 1.8 | 3.6 | 3.6 |
| ^30^N_2_ taken out | - | - | - | - | 0.0085 | 0.0028 | 0.017 | 0.0055 |
| Total N (µmol) | | | 11056.7 | 11154.0 |  | | 10536.7 | 11021.1 |
| Total N recovered at day 28 (%) | | | | | | | **95.3** | **89.8** |
